# Widespread discordance between mRNA expression, protein abundance and *de novo* lipogenesis activity in hepatocytes during the fed-starvation transition

**DOI:** 10.1101/2025.04.15.649020

**Authors:** Austin Landgraf, Junichi Okada, Maxwell Horton, Li Liu, Shoshana Solomon, Yunping Qiu, Irwin J. Kurland, Simone Sidoli, Jeffrey E. Pessin, Kosaku Shinoda

## Abstract

The mammalian liver plays a critical role in maintaining metabolic homeostasis during fasting and feeding. Liver function is further shaped by sex dimorphism and zonation of hepatocytes. To explore how these factors interact, we performed deep RNA-sequencing and label-free proteomics on periportal and pericentral hepatocytes isolated from male and female mice under fed and starved conditions. We developed a classification system to assess protein-mRNA relationship and found that gene products (mRNA or protein) for most zonation markers showed strong concordance between mRNA and protein. Although classical growth hormone regulated sex-biased gene products also exhibited concordance, ∼60% of sex-biased gene products showed protein-level enrichment without corresponding mRNA differences. In contrast, transition between feeding and starvation triggered widespread changes in mRNA expression without significantly affecting protein levels. In particular, key lipogenic mRNAs (e.g. *Acly*, *Acaca*, and *Fasn*) were dramatically induced by feeding, but their corresponding proteins (ACLY, ACC1, and FAS) showed little to no change even as functional *de novo* lipogenic activity increased ∼28-fold in the fed state. To facilitate further exploration of these findings, we developed Discorda (https://shinoda-lab.shinyapps.io/discorda/), a web database for interactive analysis. Our findings reinforce the principle that mRNA changes do not reliably predict corresponding protein levels (and vice versa), particularly in the context of sex and acute metabolic regulation of hepatocytes, and that *de novo* lipogenesis activity can be completely uncoupled from changes in protein expression.

## INTRODUCTION

Liver metabolism is tightly controlled by gene expression (including mRNA and protein levels) as well as by allosteric regulators and post-translational modifications (PTMs) of metabolic enzymes^1^. In response to metabolic stress such as starvation, hepatic gluconeogenesis is induced by increased glucagon/epinephrine and decreased insulin levels, leading to the activation of both CREB and FOXO1 transcription factors. This hormonal shift activates the cAMP-PKA pathway and inhibits Akt signaling, causing CREB phosphorylation and FOXO1 nuclear translocation, whereupon both transcription factors bind to promoters of key gluconeogenic genes (such as *Pck1* and *G6pc*)^2–4^. On the contrary, *de novo* lipogenesis (DNL) in the liver is upregulated in the fed state, primarily through the activation of the transcription factors sterol regulatory element-binding protein 1c (SREBP-1c) and carbohydrate response element-binding protein (ChREBP). Insulin activates transcription of *Srebf1* (encodes SREBP-1c) via liver X receptors (LXRs)^5^, while increased carbohydrate-derived metabolites activate ChREBP, leading to the upregulation of key lipogenic enzymes such as fatty acid synthase (FAS) and acetyl-CoA carboxylase 1 (ACC1)^6–8^.

Biological sex is also a critical determinant of liver function and disease susceptibility^9^. The liver displays marked sexual dimorphism in both physiological (i.e., drug metabolism)^10^ and pathological states (i.e., metabolic dysfunction-associated liver disease (MASLD) and hepatocellular carcinoma (HCC))^11^. One critical factor underlying this sexual dimorphism is the sex-biased expression of a large number of genes, as demonstrated in mouse^12^ and human livers^13^. Of particular significance is the role of STAT5b, a critical transcription factor in driving expression of numerous sex-biased hepatic genes. Growth hormone secreted from the anterior pituitary binds the growth hormone receptor (GHR) in the liver, causing phosphorylation of JAK2 which then phosphorylates STAT5b. In mice, GH secretion patterns are sex dimorphic with males showing pulsatile secretions and females showing continuous secretion, leading to differential activation of STAT5b and its targets^14^. Humans also exhibit sex-dimorphic GH secretion patterns but the extent of the role of STAT5b in regulating human sex-biased hepatic gene expression is not fully elucidated^15^.

In addition to metabolic state and sex, liver function is also dependent upon the spatial organization of hepatocytes, which are organized in repetitive hexagonal arrays termed lobules^16^. Hepatocytes located closer to the portal vein display increased β-oxidation, ammonia detoxification and gluconeogenesis compared to hepatocytes located near the central vein^16^. In contrast, pericentral localized hepatocytes display increased rates of glycolysis, *de novo* lipogenesis, bile acid and glutamine biosynthesis^16^. While many studies have examined the effect of metabolic status, sex, and spatial zonation of hepatic gene expression separately, an integrated analysis with both mRNA and protein level information has yet to be performed and would be highly valuable for liver researchers.

Studies conducted in the early 2000s with DNA microarray and proteomics using various model systems (yeast^17–19^, embryonic stem cell^20^, fibroblasts^21^) generally agreed that correlations between global mRNA and protein measurements are weak-to-moderate (Pearson’s *R* of 0.24^17^– 0.64^21^, explaining ≤40% of the variance). Nonetheless, the vast majority of previous omics studies on hepatic gene expression, including our own work^22,23^, have been at the mRNA level (microarray or RNA-seq)^22–27^. The generally accepted view is that increases in mRNA for stress and nutrient-responsive genes is succeeded by a coordinated increase in corresponding proteins and subsequent adaptive increase in functional activity. Recently, combining proteomics and RNA-seq, it was argued that mRNA-level changes in response to exercise cannot serve as an accurate proxy for subsequent changes in protein levels or function in skeletal muscle^28,29^. In any case, the correlation between the transcriptome and proteome throughout the liver lobule during the transition from fed to starvation states and its sexual dimorphism are important since progress in understanding the bases for metabolic pathophysiology is not possible without first understanding normal physiologic regulation.

Here, using deep RNA-sequencing and label-free mass-spectrometry-based proteomics, we have undertaken a systematic analysis of the concordance and discordance between mRNA expression and protein abundance of 5,494 gene products (GPs, mRNA or protein) in parallel with functional analysis of *de novo* lipogenic activity in the fed and starved states. We developed a classification system to assess the relationship between mRNA and protein changes within individual GPs, allowing us to distinguish concordant GPs (where mRNA and protein changes align) from discordant GPs (where they do not). Most zonation GPs showed strong concordance between mRNA and protein expression. Classical GH-regulated sex-biased GPs also exhibited high concordance, while a subset of sex-biased GPs showed protein-level bias without corresponding mRNA differences. These discordant changes appeared to be independent of GH-STAT5b signaling. Metabolic state transitions triggered widespread changes in mRNA expression of lipogenic genes, alongside robust increase in functional *de novo* lipogenesis. However, key lipogenic enzymes—such as acetyl-CoA carboxylase 1 (ACC1), ATP citrate lyase (ACLY), and fatty acid synthase (FAS)— showed little to no corresponding change at the protein level, despite the significant mRNA regulation. Our results demonstrate a novel framework for systematically analyzing mRNA-protein correlations, applicable to diverse biological contexts, and provides a useful data resource for liver researchers.

## RESULTS

### Building a dual-omics dataset of the zonal hepatocyte fed-starved response in both sexes

To investigate the regulation of liver gene expression in the normal physiologic transitions between the fed and starvation states, we established a paradigm in which mice rapidly consume food and sucrose water for 4 hours to create a uniform, fully-fed condition (**Fig. 1A**). Food and sucrose are removed after 4 hours of feeding, and the mice synchronously transition to the fasted and then starvation states. This paradigm is accomplished by first conditioning the mice to a daylight time-restricted feeding protocol on standard low-fat laboratory chow as previously described^30,31^. Mice are nocturnal animals that consume 75-90% of their total calories at night, yet our recent paper found no marked differences in the glucagon/insulin ratio, blood glucose levels, lipogenic and gluconeogenic genes between mice that were subjected to nighttime-restricted feeding and those that were subjected to daytime-restricted feeding^32^. We defined the fully fed timepoint as “0h” and the starved timepoint (24h after food removal) as “24h”.

**Fig. 1.**
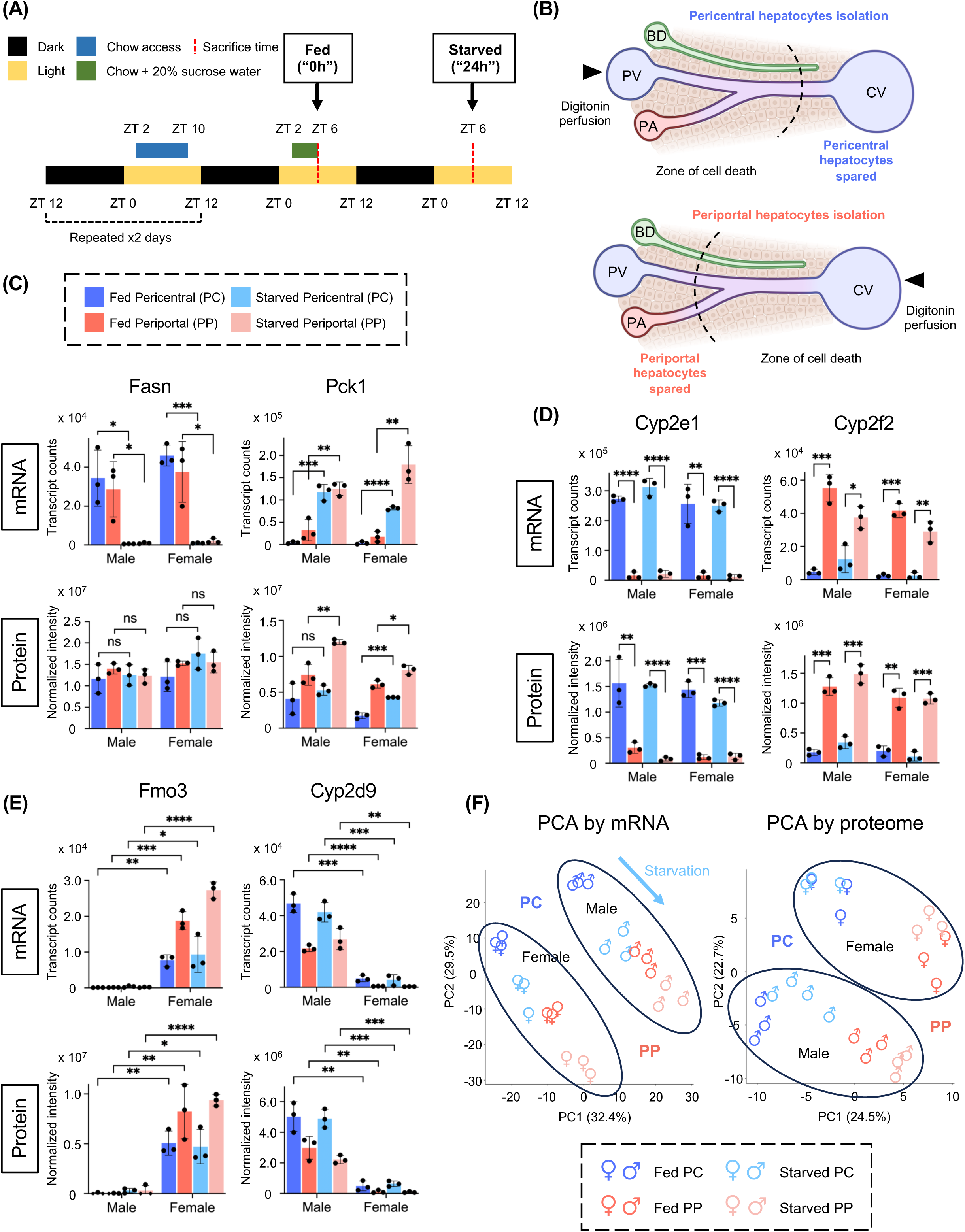
Building a dual-omics dataset of the zonal hepatocyte fed-starved response in both sexes: **(A)** Detailed scheme of the feeding and starvation protocol with zeitgeber times (ZT) indicated based on light-dark cycle. **(B)** Illustration of the digitonin ablation method in the hepatic lobule showing direction of digitonin perfusion with portal vein (PV), central vein (CV), portal artery (PA), and bile duct (BD) labeled. Lobule image adapted from BioRender. **(C-E)** RNA-seq transcript counts (upper panels) and proteomic intensity values (lower panels) for **(C)** the gluconeogenic GP *Pck1* and the lipogenic GP *Fasn*, **(D)** representative pericentral (*Cyp2e1*) and periportal (*Cyp2f2*) GPs, and **(E)** female-biased (*Fmo3*) and male-biased (*Cyp2d9*) GPs in isolated pericentral (PC) and periportal (PP) hepatocytes from male and female mice in the fed and starved states (*n*=3 mice per condition). Statistical comparisons are based on unpaired Student’s t-test performed in Prism. (n.s. = not significant, *p<0.05, **p<0.01, ***p<0.001, ****p<0.0001). Points shown as diamonds represent imputed proteomic values. **(F)** Principal component analysis (PCA) of RNA-seq and proteomics data. PCAs are based on the top 3% most variable GPs in respective datasets. Male and female samples are circled to highlight separation. PCA points represent biological replicates (*n*=3) for all 8 conditions based on coloring shown in the legend.

To assess the spatial regulation of gene expression in hepatocytes, we utilized the digitonin ablation method^33,34^ to isolate periportal (PP) and pericentral (PC) hepatocytes from male and female mice in the fed and starved states (**Fig. 1B**). All isolated hepatocytes were subjected to bulk RNA-seq and proteome-wide label-free quantification by Trapped Ion-Mobility Time-of-Flight (timsTOF HT, Bruker) Mass Spectrometry (see Materials and Methods). Independent biological replicates (*n* = 3) were used for both assays. To confirm the effectiveness of our feeding protocol, we assessed the relative induction of the lipogenic mRNA *Fasn* and the gluconeogenic mRNA *Pck1* in the RNA-seq (**Fig. 1C upper panels**). At the fed time point, *Pck1* mRNA is significantly reduced while *Fasn* mRNA is significantly increased, as expected. Twenty-four hours following the removal of food and sucrose water, *Fasn* mRNA is substantially suppressed and *Pck1* mRNA is increased, also as expected.

Whether or not changes in mRNA are reflected at the protein level is gene-specific. Here, we observed that relative changes in *Pck1* mRNA roughly correlate with its protein PEPCK in our mass spectrometry data, consistent with our recent preprint^32^ (**Fig. 1C lower panels**). However, despite the dramatic change in *Fasn* mRNA, its protein, FAS, showed no significant changes between the fed and starved states. Next, the relative purity of the isolated PP and PC hepatocytes was confirmed by assessing the mRNA and protein expression of the known zonation markers *Cyp2e1* and *Cyp2f2*. On average, *Cyp2e1* exhibited 17-fold enrichment at the mRNA level and 12-fold enrichment at the protein level in the isolated pericentral hepatocytes. Conversely, *Cyp2f2* showed 11-fold enrichment at the mRNA level and 7-fold enrichment at the protein level in the isolated periportal hepatocytes (**Fig. 1D**). Finally, we confirmed sex specific gene expression by assessing the known GH-target genes, *Fmo3* (female-biased) and *Cyp2d9* (male-biased). *Fmo3* exhibited 112-fold enrichment at the mRNA level and 124-fold enrichment at the protein level in the female hepatocytes. Conversely, *Cyp2d9* showed 33-fold enrichment at the mRNA level and 17-fold enrichment at the protein level in the male hepatocytes (**Fig. 1E**).

As shown in **Fig. 1F**, Principal Component Analysis (PCA) was applied to the RNA-seq and proteome data. In both transcriptome and proteome PCAs, male and female hepatocytes were clearly separated into distinct clusters (outlined by circles). Periportal and pericentral hepatocytes were also clearly delineated in the proteome and transcriptome PCAs. The effect of starvation and feeding was represented in the third Principal Component (PC3) (**Suppl. Fig. 1A-B**). PC3 accounted for 22.7% of data variation in the transcriptome and separated the 12 fed samples (top circle) from the 12 starved samples (bottom circle) in the mRNA PCA. In contrast, PC3 only accounted for 6.7% of data variation in the proteome and did not clearly delineate any conditions in the proteome PCA.

For an integrated analysis of RNA-seq and proteome datasets, first we directly mapped UniProt accessions to corresponding Ensembl gene IDs using UniProt ID Mapper^35^, resulting in 91.0% matches between the two datasets. While comprehensive, we wanted to minimize the number of unmatched accessions presumably due to the known discontinuity of Ensembl IDs between genome assemblies^36^. To achieve significantly higher match rate, we devised a pipeline shown in **Suppl. Fig. 1C**: (1) UniProt accession were mapped to gene symbols using UniProt ID Mapper. (2) Ensembl gene IDs were also mapped to gene symbol using “mygene” R package. (3) The two datasets were integrated at the level of gene symbols (96.7% match rate). (4) For protein IDs only found in the mass spectrometry data, UniProt accessions were directly mapped to Ensemble gene ID. With this strategy, we achieved a final 98.2% match rate between the two datasets, meaning 5,494 out of 5,592 identified proteins have corresponding gene-level mRNA quantification. This is, to the best of our knowledge, the highest coverage dual-omics dataset that describes the effects of feeding and starvation in hepatocytes. This unique dual-omics dataset enables deep investigation of three essential parameters of hepatocyte physiology: zonation, sex, and metabolic state. In the following sections, the effects of each parameter are discussed in detail.

### Two complementary systems identify parameter-specific mRNA-protein discordance

As defined by Selbach et al., two types of mRNA-protein correlations exist: *within-gene* and *across-gene* (**Suppl. Fig. 2A**)^37^. Within-gene correlations examine how mRNA changes between conditions explain corresponding protein changes. Across-gene correlations assess how mRNA and protein abundances align across different genes in a given sample. These correlations are stronger than within-gene correlations despite a vast dynamic range (∼10⁸) in mRNA and protein levels. In our dataset, across gene correlation also has a higher correlation (Pearson *R* = 0.535) than within-gene correlation (Pearson *R* = 0.095 – 0.491), (**Suppl. Fig. 2B**). A subset of genes, however, showed poor correlation between mRNA and protein abundance, such as *Alb, Apob*, *Apoa2*, *C3, Serpina1b,* and *Selenop*, all of which show disproportionally high mRNA expression (**Suppl. Fig. 2C,** outlined by a green circle). Interestingly, these genes encoded proteins that are secreted by hepatocytes into the circulation. Conversely, some mitochondrial genes (e.g. *Ndufa9* and *Ndufa4*, outlined by orange circle) showed disproportionally high protein expression. *Cps1* was the most abundant protein while *Alb* was the most abundant mRNA in our datasets, which is consistent with previous reports^38^.

Harnessing the unique dual-omics dataset, we analyzed within-gene correlations for three key biological parameters: zone (pericentral *vs* periportal), sex (female *vs* male), and metabolic state (fed *vs* starved). As shown in **Fig. 2A**, for every gene product (GP) shared between the proteome and RNA-seq datasets, we plotted the *protein* log_2_ fold change with respect to zone (PC over PP) on the y-axis versus the *mRNA* log_2_ fold change on the x-axis. We performed a linear regression (blue line) and calculated the Pearson correlation coefficient (*R*) between protein and mRNA for the zone parameter in female fed hepatocytes (*R* = 0.504). We repeated this analysis for the male fed condition which showed a similar *R* value (0.469) (**Suppl. Fig. 2D**). On average, within gene correlations for the zone parameter is strong (mean *R* = 0.491, %CV = 4.7%) across all four conditions (i.e., female fed, female starved, male fed, and male starved). The low %CV suggests the within-gene correlation for the zone parameter is consistent across sex and metabolic state. We also calculated the Spearman’s rank correlation coefficient (ρ) for all four zone conditions (mean ρ = 0.349) as shown in **Suppl. Fig. 2B (right)**, which closely approximated within-gene Spearman’s correlation for zonation in male mice (ρ = 0.39)^38^ previously reported by Itzkovitz et al..

**Fig. 2.**
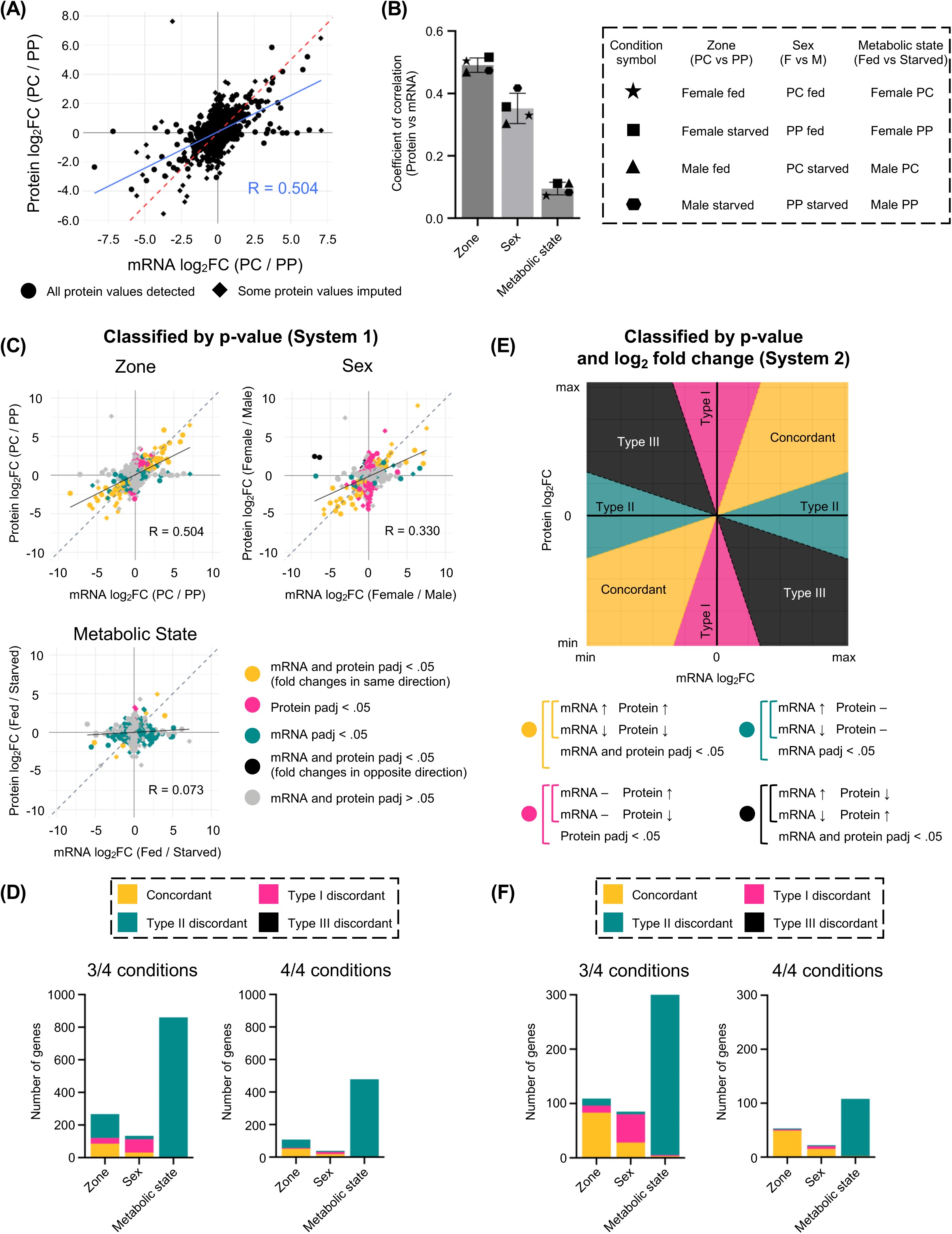
Two complementary systems identify parameter-specific mRNA-protein discordance: **(A)** Scatter plot of *within*-gene correlation between protein and mRNA log_2_ fold change (FC) for pericentral (PC) vs periportal (PP) comparison in fed female mice. Dotted red line represents slope of 1. Solid blue line shows linear fit. Pearson correlation coefficient (*R*) is shown in blue. GPs for which one or more protein values has been imputed are indicated by diamonds; for RNA-seq, no imputation was performed. **(B)** Pearson correlation coefficients (*R*) for all *within*-gene correlations plotted by biological parameter (zone, sex, and metabolic state). Each point shows the *R* value for a specific condition represented in the table by distinct shapes. **(C)** Scatter plot of *within*-gene correlation between protein and mRNA log_2_ fold change (FC) for representative pericentral (PC) vs periportal (PP) comparison (top left), female vs male comparison (top right), and fed vs starved comparison (bottom left) with points colored by adjusted p-value as indicated in bottom right panel. Dotted line represents slope of 1, and solid black line represents linear fit with Pearson *R* shown. Conditions shown: zone = female fed, sex = PC fed, metabolism = female PC. **(D)** The number of GPs belonging to the concordant (gold), type I (pink), type II (green), and type III (black) discordant categories as defined in (C) according to adjusted p-value (system 1). 3/4 = classified in the corresponding category in three or more of the four conditions. 4/4 = all four conditions. **(E)** Visual description of system which assigns GPs to four categories based on adjusted p-value and log_2_ fold change. Image created using Desmos. **(F)** The number of GPs belonging to the concordant (gold), type I (pink), type II (green), and type III (black) discordant categories as defined by system 2. 3/4 = classified in the corresponding category in three or more of the four conditions. 4/4 = all four conditions.

Next, we compared the mean *R* for each *parameter* (zonation, sex, and metabolic state) across all *conditions* (**Fig. 2B**). Each *parameter* was analyzed across *four conditions* which are represented by distinct symbols shown in the adjacent box. The three parameters showed distinct levels of correlation with zone showing the highest (mean *R* = 0.491, %CV = 4.7%), then sex (mean *R* = 0.353, %CV = 13.8%), and finally metabolic state (mean *R* = 0.095, %CV = 21.7%). The low mean *R* for the metabolic state parameter suggested the acute mRNA response to feeding shows virtually no correlation with protein changes (0.095^2^ = ∼0.9% of the variability in protein levels can be explained by mRNA changes).

While the aforementioned approach is useful to assess the overall strength of correlations, it is essential to determine the mRNA-protein relationship for individual gene products (GPs). We aimed to systematically identify GPs with a strong mRNA-protein correlation (concordant). Conversely, for GPs with poor mRNA-protein correlation (discordant), their discordance can be classified into two types: (1) protein changing but not mRNA (2) mRNA changing but not protein. We devised two methods to classify GPs accordingly. The first method, which we named system 1 (**Fig. 2C and 2D**), classified GPs for zone, sex and metabolic state into four categories based on Benjamini–Hochberg adjusted p-value. These categories were: (1) protein adjusted p-value < 0.05 & mRNA adjusted p-value < 0.05 with fold changes in the *same* direction (concordant – colored gold) (2) protein adjusted p-value < 0.05 & mRNA adjusted p-value > 0.05 (designated as type I discordant – colored pink) (3) protein adjusted p-value > 0.05 & mRNA adjusted p-value < 0.05 (type II discordant – colored green) (4) protein adjusted p-value < 0.05 & mRNA adjusted p-value < 0.05 with fold changes in *opposite* directions (type III discordant – colored black). Representative examples of the system1-based classification are shown in **Fig. 2C**. Concordant GPs (gold) were abundant in the sex and zone comparisons. Type I discordant GPs (pink) were most abundant in the sex comparison, and type II discordant GPs (green) were the major category found in the metabolic state comparison. Next, we quantified the number of GPs in each category (concordant, type I, II, and III discordant) (**Fig. 2D**). Importantly, classification was not always consistent across our four conditions because certain GPs exhibit sex-specific zonal expression and zone-specific metabolic regulation^39,40^. In our analysis, to be defined as concordant or discordant, a GP had to be classified as such in three out of four (3/4) conditions (**Fig. 2D**). In system 1 at 3/4 cutoff, zone parameter was comprised of roughly 55% type II, 32% concordant, and 13% type I GPs. The sex parameter consisted of 61% type I, 23% concordant, and 16% type II GPs. The metabolic state parameter was almost entirely (>99%) type II discordant GPs, indicating that despite having the highest number of significantly changed GPs among the three parameters, the metabolic state-induced changes at the mRNA level were hardly reflected at the protein level during the 24-hour period. Although some GPs showed type III discordance in one of the conditions, none reached the 3/4 condition cutoff. The trends described above were consistent whether we defined GPs based on 3/4 or 4/4 cutoffs. The number of GPs in each category at all cutoffs (1/4, 2/4, 3/4, and 4/4 conditions) is shown in **Suppl. Fig. 2E**.

While the p-value-based system 1 is useful, it is not sufficiently strict. For example, a gene product that displays 1.2-fold enrichment at the protein level and a 24-fold enrichment at the mRNA level can still be classified as concordant despite a 20-fold difference in degree of enrichment *if* both enrichments are statistically significant. To address this, we used a more stringent method, we named system 2, that classifies GPs based on both p-value and fold change (**Fig. 2E and 2F**). Briefly, the protein vs mRNA scatter plot is divided into 8 regions by straight lines y = 3x and y = 1/3x as shown in **Fig. 2E**, where x represent mRNA log_2_ fold change and y represents protein log_2_ fold change. The slopes of these lines stipulate that the relative protein and mRNA log_2_ fold change must be within a factor of 3-fold for a GP to be classified as concordant. GPs were classified based on the shaded region in which they fall: Gold as concordant, pink as type I discordant, green as type II discordant, and black as type III discordant. Similar to the first system, GPs must also meet appropriate adjusted p-value cutoffs. This approach has the advantage of conservatively estimating the number of discordant GPs over the p-value only based system 1. As shown in **Fig. 2F**, system 2 showed very similar trends as system 1 with one exception: the zone parameter was comprised of 76% concordant GPs as opposed to 32% in system 1 at the 3/4 cutoff. Consistent with system 1, system 2 showed that the sex parameter contained the highest percentage (also 61%) of type I GPs. In system 2, the metabolic state parameter was again almost entirely type II discordant GPs (98%). As with the system 1, at the 3/4 cutoff, no GPs showed type III discordance. Together, these two complementary algorithms demonstrate that the degree of mRNA-protein correlation is highly parameter-dependent. Zonal gene expression showed the strongest concordance, while the sex parameter exhibited a significant number of type I discordant GPs. To our surprise, metabolic state-induced changes exhibited widespread discordance that suggested protein levels are decoupled from mRNA changes during the early metabolic response. The number of GPs in each category based on system 2 at all cutoffs is shown in **Suppl. Fig. 2F**.

Because there is no agreed upon method to define concordant and discordant GPs, we opted to be conservative with our classification and used the more stringent system 2 for all subsequent figures.

### Systematic identification of zonal markers concordant between mRNA and protein

With the well-recognized advent of single-cell RNA-seq and the recent development of low-input spatial proteomics^41^, there is an increasing need to identify zonation markers that are concordant between mRNA and protein. In a seminal paper, Itzkovitz’s group used surface antigens and FACS to collect hepatocytes from multiple zones. Subsequent RNA-seq and LC-MS/MS analysis showed that mean mRNA and protein expressions across the lobule were well correlated^38^. Naef’s group examined the effects of circadian rhythms to identify circadian-independent zonation markers using scRNA-seq^42^. Here, we used an orthogonal antibody-independent approach to physically isolate pericentral and periportal hepatocytes in both sexes under well-controlled fed and starved conditions. This allowed us to identify robust markers that are unaffected by metabolic state or sex. As shown in **Fig. 3A**, in the female fed state, system-2-based screening identified numerous concordant markers including the well-known *Glul* and *Gulo* with enrichment in PC hepatocytes, and *Cyp2f2* and *Cdh1* with enrichment in PP hepatocytes. We applied the same system 2 algorithm to the remaining three conditions of PC/PP hepatocytes and overlapped concordant GP sets. As shown by UpSet plot in **Fig. 3B**, 18 periportal and 31 pericentral markers (total 49 GPs) were concordant under all conditions. These 49 highly mRNA-protein concordant markers exhibited zonation independent of sex or the metabolic states examined and are shown as a heatmap in **Fig. 3C**. These included the well-known *Cyp2e1* (PC), *Cyp2f2* (PP), and *Oat* (PC), as well as less familiar markers, such as *Aldh1b1* and *Cryl1*. Transcript and protein intensities of representative sex and metabolic-state independent markers with biological replicates are shown in **Fig. 3D** (pericentral GPs) and **Fig. 3E** (periportal GPs). These GPs for portal and central hepatocytes can be used as useful markers for future exploration of liver zonation.

**Fig. 3.**
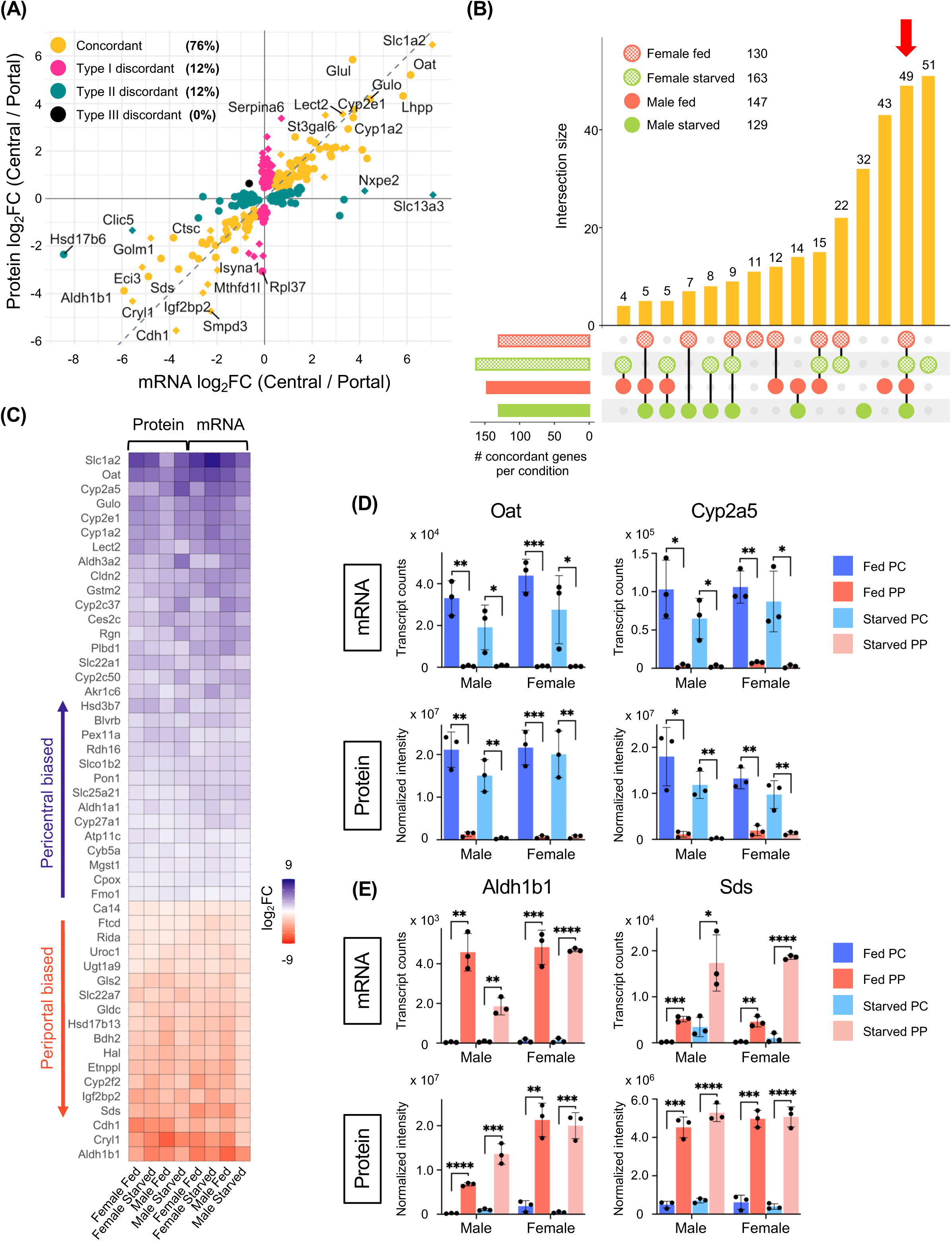
Systematic identification of zonal markers concordant between mRNA and protein: **(A)** Scatter plot with coloring based on system 2 of protein vs mRNA log_2_ fold change for pericentral (PC) vs periportal (PP) comparison in fed female mice. Dotted gray line represents slope of 1. Diamonds represent GPs with imputed protein values. Percentages based on 3/4 condition cutoff. **(B)** UpSet plot of concordant zonation GPs across female fed, female starved, male fed, and male starved conditions with set sizes indicated. Gene products showing mRNA-protein concordance in all four conditions (49 GPs) are indicated by the red arrow. **(C)** Protein and mRNA expression heatmap of the 49 zonation markers from (B) with columns representing individual conditions. Color scale is based on log_2_ fold change (PC/PP). **(D-E)** RNA-seq and proteomic values for representative sex and metabolic state independent **(D)** pericentral and **(E)** periportal markers (*n*=3 mice per condition). Statistical comparisons are based on unpaired Student’s t-test performed in Prism. (n.s. = not significant, *p<0.05, **p<0.01, ***p<0.001, ****p<0.0001).

### A sex-biased expression observed exclusively at the protein level and its potential regulators

The liver exhibits sex dimorphism in both physiological and pathological states^10,11^. Partially underlying these phenotypic differences is a sexual dimorphic expression of a large number of genes. STAT5b emerges as a critically important factor in driving sex-biased expression of many hepatic genes, reflecting its established role as a master regulator of sex differences through the male-specific pulsatile growth hormone (GH) signaling^43–45^. As shown in **Fig. 4A**, system 2-based screening found that sex dimorphic GPs were 33% concordant, 61% type I discordant, and 6% type II discordant. In the fed pericentral hepatocytes, many previously reported sex dimorphic GPs, such as *Fmo3*, *Hao2* (female-biased) and *Cyp4a12a*, *Mup1* (male-biased) exhibited concordance. We also identified GPs that only showed sexual dimorphic expression at the protein level (type I discordance), most of which have not been previously reported. The comprehensive list of sex-biased GPs at the 3/4 cutoff (28 concordant and 51 type I) is shown as a heatmap in **Fig. 4B**. Repeatedly concordant sex GPs included *Sult2a1*, *Acot3* (female-biased), *Hsd3b5,* and *Selenbp2* (male-biased). Type I discordant sex GPs we identified included *Hmgb2*, *Saraf, H2-D1* (female-biased), *Gpx3*, *Atp6v0d1*, and *Akr1c13* (male-biased). Of note, with the exception of *Eif2s3y* (Y-chromosome), *Fmr1* (X-chromosome), and *Timm8a1* (X-chromosome), all concordant and type I sex GPs shown are coded on autosomal chromosomes.

**Fig. 4.**
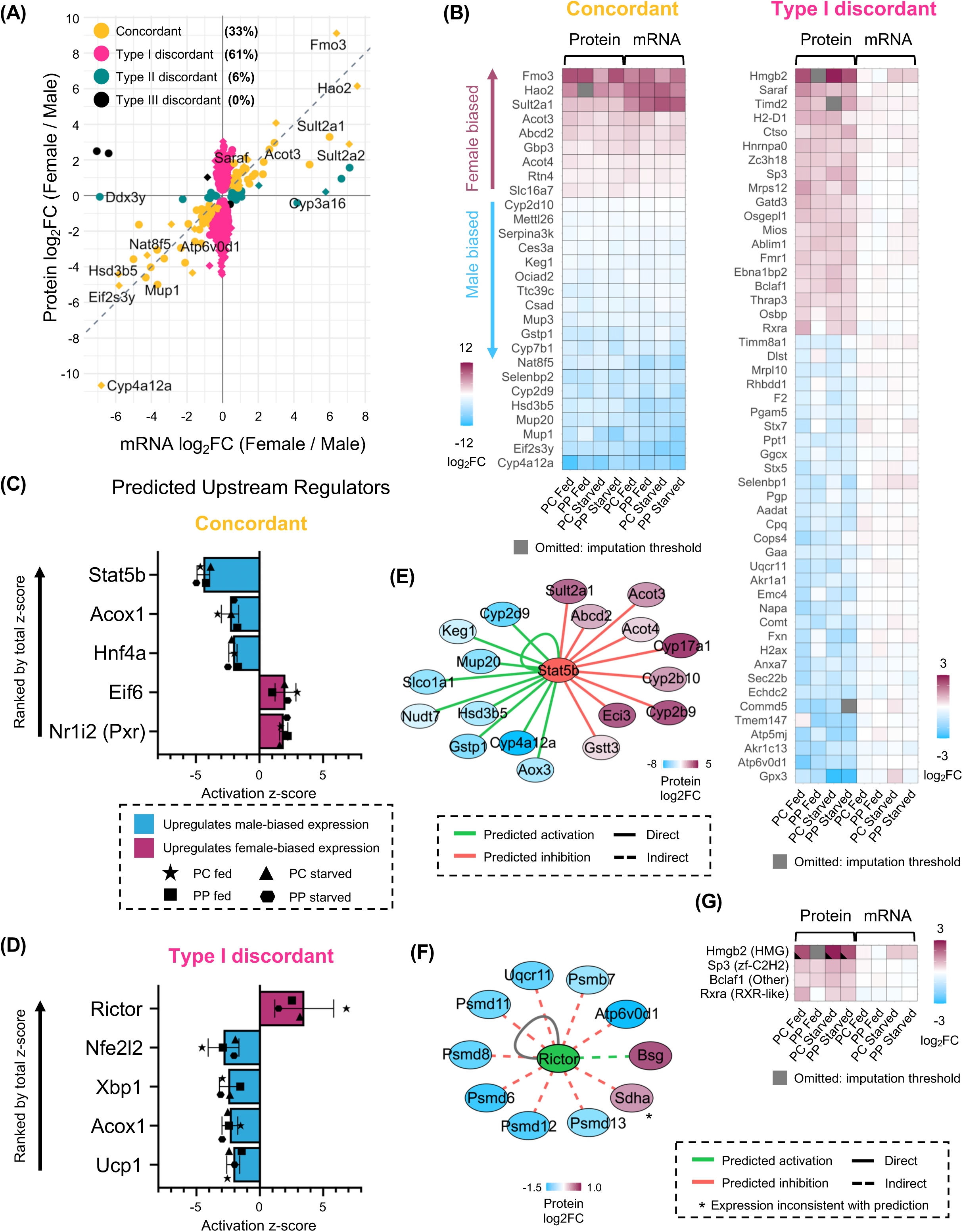
A sex-biased expression observed exclusively at the protein level and its potential regulators: **(A)** Scatter plot with coloring based on system 2 of protein vs mRNA log_2_ fold change for female vs male comparison in fed pericentral hepatocytes. Dotted gray line represents slope of 1. Diamonds represent GPs with imputed protein values. Percentages based on 3/4 condition cutoff. **(B)** Protein and mRNA expression heatmap of 28 concordant (left panel) and 51 type I discordant (right panel) sex-biased GPs with columns representing individual conditions. Color scale is based on log_2_ fold change (female/male). Only GPs classified as concordant / type I discordant in 3/4 or 4/4 conditions are shown. Protein data shown in gray were excluded from the analysis because they did not meet the criteria for imputation using the MinDet algorithm. **(C-D)** Qiagen IPA Upstream Regulator Analysis of the concordant and type I discordant GPs in each condition based on protein log_2_ fold change (female/male). Upstream regulators are ranked based on their activation z-score. Plots were generated by performing Comparison Analysis of individual Core Analysis results for each condition. Z-scores for individual conditions are represented by distinct shapes as shown in the legend. **(E-F)** Top predicted upstream regulators and their downstream targets (from IPA) for concordant and type I discordant GPs as shown in Cytoscape. For STAT5b **(E)**, predictions and observations agree for all 18 GPs. For RICTOR **(F)**, predictions and observations match for 9 out of 10 GPs. The average protein log_2_ fold change (*n* = 3) is shown for each target. Regulatory networks shown are based on the fed PP condition. **(G)** Protein and mRNA expression heatmap of known transcription factors from the AnimalTFDB database showing protein and mRNA log_2_ fold change (female / male) for each condition. Transcription factor family is shown in parentheses. Only transcription factors classified as concordant or discordant in 3/4 or 4/4 conditions are shown. Black wedges represent proteins with imputed values. Proteins shown in gray were excluded from the analysis because they did not meet the criteria for imputation using the MinDet algorithm.

Interestingly, applying 4/4 cutoff resulted in only 15 concordant (i.e., *Sult2a1, Abcd2, Cyp4a12a, Hsd3b5*) and 5 type I discordant GPs (*Atp6v0d1, Fmr1, Ablim1, Bclaf1, Thrap3*). We postulated that the large discrepancy in the number of concordant and type I GPs between 3/4 and 4/4 cutoff was due to zone-specific sex-bias, which has been previously reported^46^. We quantified the number of zone-specific concordant and type I discordant sex GPs by UpSet plot (**Suppl. Fig. 3A and B**). For the concordant sets, there were 13 periportal-specific and 12 pericentral specific sex GPs. For the type I sets, we identified 5 periportal-specific and 49 pericentral specific sex GPs. These GPs are shown in the form of heatmaps in **Suppl. Fig. 3C-D** for concordant and **Suppl. Fig. 3E-F** for the type I set. Representative periportal sex-biased concordant GPs such as *Cd36* (fatty acid translocase), and *Adh4* (alcohol dehydrogenase 4), are plotted in **Suppl. Fig. 3G.**

Next, we explored possible regulators of concordant and type I discordant sex GPs using Upstream Regulator (UR) Analysis (**Fig. 4C-D**)^47^. The top UR predicted for concordant GPs was STAT5b, consistent with reports by Waxman’s group and previous studies^44^. Of note, STAT5b was predicted as the top UR of concordant GPs regardless of whether mRNA or protein expression was used as the input. HNF4A, which has previously been identified as a regulator of sex dimorphic gene expression, was also among the top predicted UR^48^. In contrast, STAT5b was not a top UR for the type I discordant GPs (**Fig. 4D**). In this category, top predicted regulators included RICTOR (a component of mTORC2 that influences protein translation and stability) and NFE2L2 (also known as NRF2). Depletion of RICTOR has been shown to significantly decreases male lifespan but not female lifespan in mice, although the mechanism behind this is not fully understood^49^. Likewise, NRF2 activation in the liver is associated with sexual dimorphism^50^. The regulatory networks for top upstream regulators of concordant (STAT5b) and type I discordant GPs (RICTOR) are shown in **Fig. 4E and 4F** respectively with their downstream targets colored by fold change.

Historically, transcription factors (TFs) have been difficult to detect in proteome datasets due to their low abundance. Because of the high coverage and sensitivity of the timsTOF HT mass spectrometer coupled online with nano LC, we were able to identify 150 TFs as annotated in AnimalTFDB^51^. We screened our concordant, type I, and type II GPs sets against these 150 TFs and identified 4 transcription factors which showed sex bias in 3 or more out of 4 conditions (**Fig. 4G**). These consisted of *Hmgb2*, *Sp3*, *Bclaf1* and previously reported *Rxra* (female-biased)^52^. Interestingly, all of these TFs showed type I discordance meaning that transcriptome analysis alone would have missed their sex dimorphism. Despite being a top upstream regulator, there was no significant difference between males and females in either STAT5b mRNA or protein, which is consistent with the known activation mechanism of STAT5b through phosphorylation by Janus kinase 2 (JAK2).

In summary, we found two types of sex dimorphism in hepatocyte gene expression: classical STAT5b targets which mostly exhibit concordance and previously unreported protein-only sex-biased gene products which do not appear to be regulated by STAT5b. This data illustrates that sex-biased gene regulation in mouse hepatocytes can arise from distinct mechanisms of control.

### Widespread mRNA-protein discordance induced by acute feeding and starvation

The third parameter in our dual-omics dataset is metabolic state (fed vs starved). We applied system 2 based screening and surprisingly saw that 98% of GPs exhibited significant changes only at the mRNA level (type II discordance) in the two timepoints we examined (fed for 4 hours and starved for 24 hours), as shown in the female pericentral condition (**Fig. 5A**). These type II GPs include numerous enzymes in the lipogenesis pathway, such as *Fasn*, *Acly*, and *Acaca*, as well as the gluconeogenesis pathways, *Pck1* and *Sds*. At the 3/4 cutoff, there were 299 type II discordant, 2 type I discordant, and 3 concordant metabolic GPs. Although the number of GPs in the concordant category is small, this list included the critical lipogenic transcription factor, *Srebf1* (encodes SREBP-1a and SREBP-1c). The remaining two concordant metabolic GPs were previously reported allosteric regulators of lipogenesis (*Mid1ip1* – encodes MIG12) and gluconeogenesis (*Pdk4* – encodes PDK4). MIG12 was reported by Horton’s group to positively regulate fatty acid synthesis *in vivo* by inducing polymerization of ACC^53^, a critical enzyme in the lipogenesis pathway. PDK4 is a mitochondrial protein that phosphorylates the pyruvate dehydrogenase (PDH) complex, inhibiting the conversion of pyruvate to acetyl-CoA. PDK4 was reported to stimulate both gluconeogenic gene expression and fatty acid oxidation^54^. Similar to the concordant category, only two type I discordant GPs met the 3/4 cutoff: programmed cell death protein 4 (PDCD4) (starvation-induced) and transferrin (TF) (fed-induced). PDCD4 is a tumor suppressor protein that inhibits translation initiation by directly binding to the entry channel of the 40S ribosomal subunit through its N-terminal domain while using its C-terminal domain to sequester and inactivate eIF4A helicase, particularly during cellular stress conditions^55,56^. Intermittent fasting is reported by Larance et al. to induce PDCD4 protein in the mouse liver^57^, although the authors did not investigate the mRNA level change. Transferrin is an iron-binding protein produced by the liver and secreted into the blood that is important for maintaining iron homeostasis^58^. Fasting is reported to regulate plasma iron levels in mice and humans^59^, which could account for the parallel regulation of transferrin protein in the liver (Note that transferrin is represented by the gene symbol *Trf* (Ensembl) in our mRNA data and *Tf* (UniProt) in our protein data). Some GPs showed concordance in the female PC condition but did not meet the 3/4 cutoff we chose to use, as shown in **Fig. 5A**. These included *Insig1* and *Bhlhe40* (concordant, fed-induced), which are critical regulators of SREBP-1c^60,61^. Similarly, *Slc25a47*, which has been shown to regulate gluconeogenesis^62^, exhibited concordant fasting-induction in the female PC condition but also did not meet the 3/4 cutoff. This may suggest a sex-specific or zone-specific regulatory mechanism.

**Fig. 5.**
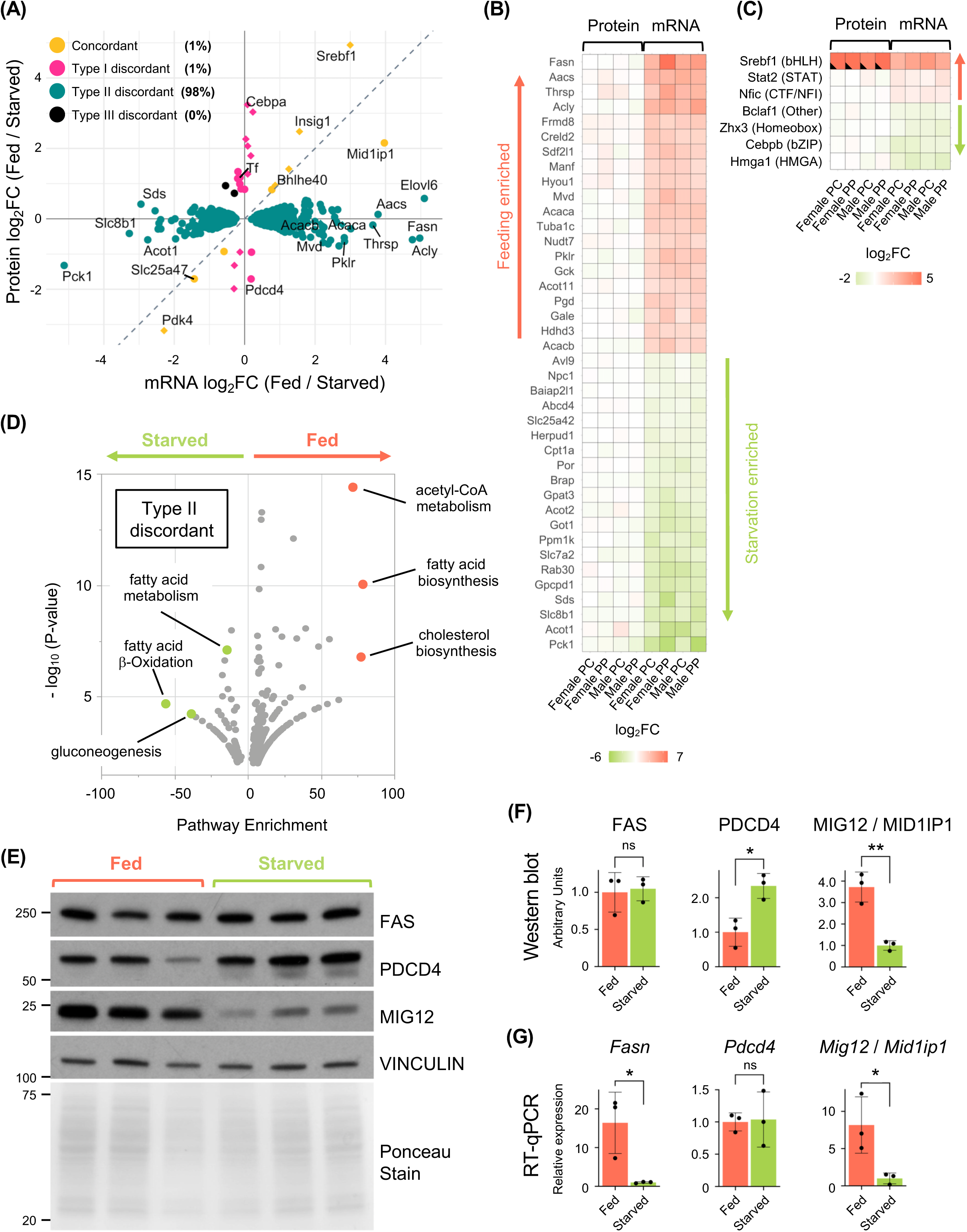
Widespread mRNA-protein discordance induced by acute feeding and starvation: **(A)** Scatter plot with coloring based on system 2 of protein vs mRNA log_2_ fold change for fed vs starved comparison in female pericentral hepatocytes. Dotted gray line represents slope of 1. Diamonds represent GPs with imputed protein values. Percentages based on 3/4 condition cutoff. **(B)** Protein and mRNA expression heatmap of the top 20 fed-induced and top 20 starvation-induced type II discordant GPs. GPs are sorted by average mRNA and protein fold change. Color scale is based on log_2_ fold change (fed/starved). Only GPs classified as type II discordant in 4/4 conditions are shown. **(C)** Protein and mRNA expression heatmap of known transcription factors from the AnimalTFDB database showing protein and mRNA log_2_ fold change (fed/starved) for each condition. Transcription factor family is shown in parentheses. Only transcription factors classified as concordant or discordant in 3/4 or 4/4 conditions are shown. Black wedges represent proteins with imputed values. **(D)** Metascape functional enrichment analysis based on mRNA fold change of all type II discordant GPs meeting 4/4 cutoff (32 starvation-induced; 74 fed-indcued) plotted as enrichment score vs negative log_10_(p-value). **(E-G) (E)** Representative immunoblotting of FASN (type II), PDCD4 (type I), MID1IP1 / MIG12 (concordant) and Vinculin (loading control) from total liver samples from fed and starved male mice. Total protein loading is shown by Ponceau staining. (**F**) Quantification of the bands in (E) normalized to Vinculin. (**G**) Results of RT-qPCR from the same samples shown in (E-F) for *Fasn, Pdcd4,* and *Mig12 / Mid1ip1* mRNA normalized to *Apoa2* mRNA. Statistical comparisons are based on unpaired Student’s t-test performed in Prism. (n.s. = not significant, *p<0.05, **p<0.01).

Because over 98% GPs were type II discordant at 3/4 cutoff, we further narrowed down the 299 type II discordant list using the more stringent 4/4 cutoff to 106 “core” GPs, which are listed in **Suppl. Table 1**. In **Fig. 5B**, we plotted the top 20 feeding- and starvation-enriched GPs as a heatmap. The core feeding induced GPs included *Aacs, Thrsp*, *Mvd* and *Pklr*. Known β-oxidation GPs, such as *Acot1* and *Cpt1a*, were induced in the starved state. Not any of these mRNAs showed corresponding changes at the protein level. We also plotted all transcription factors significantly regulated by feeding and starvation as a heatmap in **Fig. 5C**. With the exception of the nutrient-dependent TF *Srebf1* (encodes SREBP-1a / SREBP-1c), which is among the three concordant metabolic GPs, all other feeding-starvation regulated TFs displayed type II discordance. These include *Stat2* (fed-induced) and *Cebpb* (C/EBPβ) (fasting-induced), which has been shown to activate gluconeogenesis^63^. The rapid concordant change in *Srebf1* mRNA and SREBP1 protein is consistent with their known regulatory mechanisms^64^.

Type II discordance during feeding and starvation occurs not only at the individual gene product level but also at the pathway level. The core GP set is particularly enriched for metabolic processes as shown in the functional enrichment analysis volcano plot in **Fig. 5D**. Enriched pathways included lipogenesis-related pathways (acetyl-CoA metabolism, fatty acid synthesis), gluconeogenesis, and β-oxidation.

The unexpected widespread discordance and the minimal number of protein level changes in the acute response to feeding motivated us to validate our result in independent samples using alternative methods (western blot and RT-qPCR). An independent set of male mice were subjected to the same feeding protocol (**Fig. 1A**) and whole liver was collected in the fed and starved states. We chose one representative gene product for each GP category based on availability of validated antibodies: *Fasn* / FAS (type II discordant), *Pdcd4* / PDCD4 (type I discordant), and *Mid1ip1* / MIG12 (concordant). As shown in **Fig. 5E and F**, western blot showed the same regulatory pattern identified by proteomics. FAS protein showed no significant change between the starvation and fed states. MIG12 was induced approximately 3.8-fold (6-fold by proteomics) with feeding, and PDCD4 protein was induced by ∼2.3-fold (2.8-fold by proteomics) with starvation. RT-qPCR results also mirrored the RNA-seq data: *Fasn* mRNA showed 16-fold induction in the fed state (40-fold by RNA-seq). *Pdcd4* mRNA showed no significant change, and *Mid1ip1* mRNA (encodes MIG12) showed 8-fold induction in the fed state (15-fold by RNA-seq) (**Fig. 5G**). RT-qPCR results were normalized to highly abundant *Apoa2* mRNA as a housekeeping gene because its expression showed no significant difference between feeding and starvation in our RNA-seq data (**Suppl. Fig. 4A**). Overall, these results are consistent with our earlier finding that *Fasn* exhibits type II discordance, *Pdcd4* exhibits type I discordance and *Mid1ip1 /* MIG12 exhibits concordance under the metabolic states tested.

The large discrepancy between the mRNA and protein for core enzymes in the *de novo* lipogenic pathway^65^, *Acly* / ACLY (**Fig. 6A**), *Acaca* / ACC1 (**Fig. 6B**), *Fasn* / FAS (**Fig. 1C**), and *Elovl6* / ELOVL6 (**Fig. 6C**) prompted us to measure functional lipogenic activity. Using the scheme illustrated in **Fig. 6D** under the same fed and starved metabolic conditions, we assessed *de novo* lipogenic activity in whole liver from male mice by measuring incorporation of deuterated water (D_2_O) into the saturated fatty acid palmitate (C16:0), utilizing a technique previously developed by Burgess and Browning^66,67^. As expected, in the fasted state there was minimal *de novo* lipogenic activity in the liver, but following 4h of feeding, *de novo* lipogenesis was induced ∼28-fold despite the constant level of lipogenic enzymes (**Fig. 6E**). Excluding the one outlier sample with highest *de novo* palmitate synthesis, the fed-induction of *de novo* lipogenesis was still ∼22-fold. As depicted in **Fig. 6F**, mRNAs for the core lipogenic enzymes are substantially (19–40-fold) induced in the fed state while their corresponding proteins remain essentially unchanged. Nonetheless, the product of the lipogenesis pathway (palmitate) is significantly elevated during refeeding. These data clearly demonstrate that lipogenic protein levels are completely dissociated from lipogenic activity in the acute fed state, even after four hours of feeding.

**Fig. 6.**
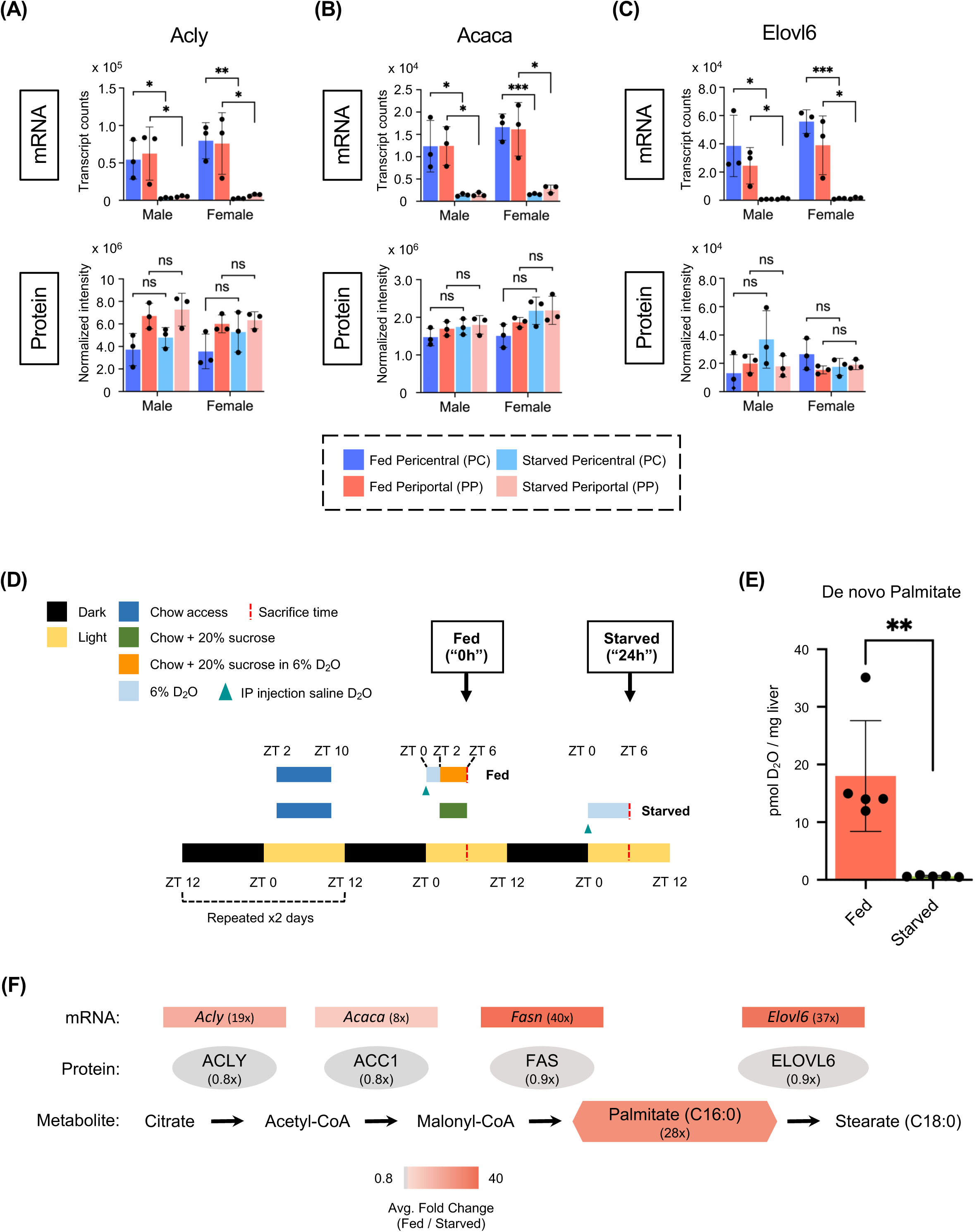
*De novo* lipogenic activity is uncoupled from changes in enzyme abundance: **(A-C)** RNA-seq transcript counts (upper panels) and proteomic intensity values (lower panels) for the core lipogenic GPs **(A)** *Acly*, **(B)** *Acaca*, and **(C)** *Elovl6* in isolated pericentral (PC) and periportal (PP) hepatocytes from male and female mice in the fed and starved states (*n*=3 mice per condition). Statistical comparisons are based on unpaired Student’s t-test performed in Prism. (n.s. = not significant, *p<0.05, **p<0.01, ***p<0.001). Points shown as diamonds represent imputed proteomic values. **(D)** Modified feeding scheme used for functional lipogenesis experiments involving deuterated water. **(E)** *De novo* palmitate measurements using deuterated water in whole liver samples from male mice in fed or starved state. Statistical comparison based on unpaired Student’s t-test performed in Prism. (**p<0.01). **(F)** Schematic illustrating the *de novo* lipogenesis pathway. Transcripts, proteins, and metabolites (palmitate) are colored based on the average Fed / Starved fold-change (non-log transformed) in RNA-seq (12-biological-replicate average), proteomics (12-biological-replicate average), and D_2_O assay (5-biological-replicate average) respectively.

### Creating the Discorda database to support the metabolic research community

To enable efficient use of this data by the scientific community, we have created a database named Discorda (https://shinoda-lab.shinyapps.io/discorda/). Discorda is equipped with four distinct visualization functions: volcano plots, heatmaps, scatter plots and classification pie charts. The volcano plot function provides a comprehensive view of mRNA and protein significance *vs* fold change across different parameters and conditions (**Fig. 7A**). This view is further enhanced by the ability to overlay a user-provided gene list and see the gene name for every data point by mouseover. Complete RNA-seq and proteome datasets are shown as independent volcano plots side-by-side for easy comparison. In addition to images, underlying RNA-seq and protein data can be downloaded in CSV format enabling users to easily conduct their own bioinformatic analysis. The heatmap function (**Fig. 7B**) displays mRNA expression and protein abundance values for user-provided gene names. Intensity values are colored based on two normalization methods which users can choose: “centered” (value – mean) and “z-score” ((value-mean)/standard-deviation). Adjacent to the gene name, our GP categories are displayed, enabling users to discern the discordance in their gene of interest across sex, zone and metabolic state. The scatter plot function (**Fig. 7C**) allows users to visualize within-gene correlation for all 5,494 mapped gene products. The categorization (concordant, discordant) identified by our two algorithms (system 1 and 2) is presented in intuitive color coding. Finally, the classification pie chart function (**Fig. 7D**) allows users to visualize absolute number and percentage of gene products belonging to each category. Like the volcano plot function, all underlying data can be downloaded in CSV format. The Discorda database will support researchers investigating hepatic sex differences, spatial arrangement, and liver metabolism by facilitating hypothesis generation, thereby advancing liver research.

**Fig. 7.**
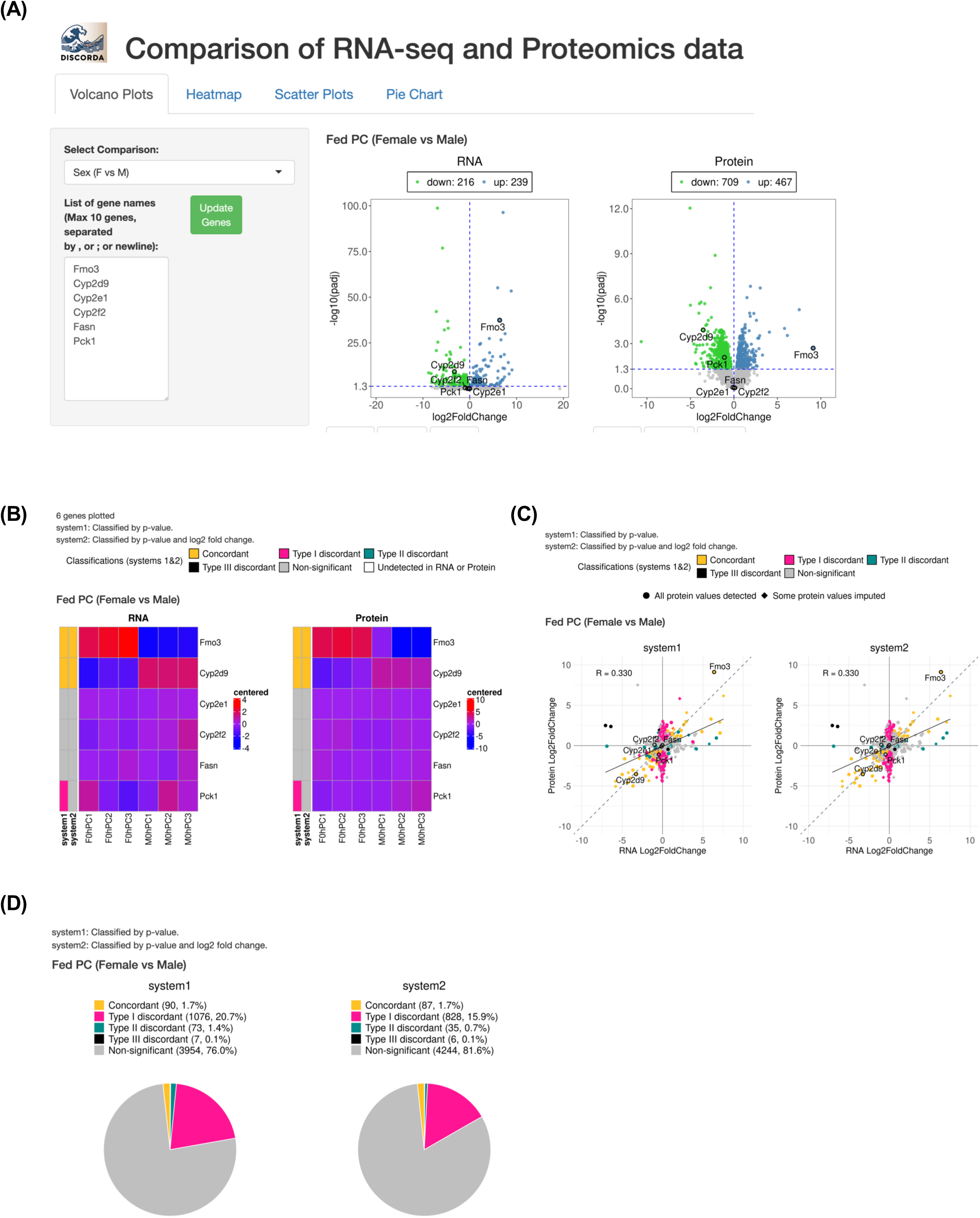
Creating the Discorda database to support the metabolic research community: **(A)** Screenshot of the Discorda landing page and the Volcano plot function. **(B)** Discorda heatmap function. **(C)** *Within*-gene protein-mRNA correlation scatter plots. **(D)** Summary pie chart function.

## DISCUSSION

Our findings build upon the body of pioneering omics study that has persisted since the 2000s (reviewed in Buccitelli & Selbach^37^), which revealed that mRNA-protein relationships can differ substantially across genes (across-gene) and within individual genes across conditions (within-gene) and further demonstrate that within-gene correlation can be dramatically shaped by biological context. Here, our comprehensive within-gene study examines three parameters that operate on different timescales: biological sex (determined in utero), zonation (postnatally shaped but somewhat plastic^32^), and metabolic state (shifting within hours). We show that (1) zonation is tightly concordant at the mRNA-protein level, (2) sex differences often manifest as protein-only biases, and (3) rapid metabolic transitions (feeding vs starvation) exhibit almost complete mRNA-protein mismatch during the early response. These findings underscore and extend the notion that transcript-based measurements alone cannot predict final protein levels and highlight how biological context and post-transcriptional regulation jointly define the mRNA-protein relationships and outcome of gene expression.

First, our data demonstrates that many zonation gene products exhibit concordance between mRNA and protein independent of sex or metabolic state. These vetted markers for portal and central hepatocytes can be used as stable landmarks for future exploration of protein zonation. Second, we identified that many GH-regulated, sex-biased gene products are concordant between mRNA and protein, but a significant subset (61%) of sex GPs are biased only at the protein level (type I discordant). RICTOR, a subunit of mTORC2, was the top upstream regulator of these type I discordant sex GPs (**Fig. 4D-F**). Functional enrichment analysis suggested many of these type I discordant sex-biased GPs (particularly male-biased GPs) are involved in the ubiquitin-proteasome system (**Suppl. Fig. 3H**). These GPs include subunits of the 20S proteasome core (*Psmb7)* and the 19S regulatory complex (*Psmd11*). mTORC1 activation broadly inhibits the ubiquitin-proteasome system (UPS)^68^, but the relationship between mTORC2 (containing RICTOR) and UPS is not well understood. Other groups suggested that mTORC2 regulates the proteasomal degradation of specific substrates (e.g. Sgk1^69^ and IRS-1^70^) through interaction with Cullin family members. It may be possible that the activation of RICTOR mediates proteasome-dependent degradation in a sex-biased manner, resulting in the type I discordance we observed.

Lastly, we found that most GPs exhibit type II discordance in response to starvation and feeding. This discordance is seen with mRNA and proteins involved in lipogenesis, gluconeogenesis, and β-oxidation. Given the high sensitivity and reproducibility of our mass spectrometry data (e.g. detection of over 100 low abundance transcription factors), it is unlikely that the lack of protein level changes is due to low assay sensitivity. Additionally, type II discordance classification does not correlate with protein abundance (**Suppl. Fig. 4B**). In general, proteins at low concentrations exhibit low signal-to-noise ratios due to the limited number of peptides detected, resulting in reduced precision and accuracy in protein quantification^71^. However, our type II discordant GPs were found in the high, medium, and low abundance range for proteins. Consequently, the absence of significant differences at the protein level cannot be attributed to measurement error. Furthermore, immunoblotting and RT-qPCR of the type I discordant gene product *Pdcd4*, the type II discordant gene product *Fasn*, and the concordant gene product *Mig12* confirmed the proteomic analyses (**Fig. 5E-G**).

Most importantly, despite the lack of protein changes, we observed a 28-fold change in functional lipogenesis. Several studies have reported the effect of fasting and feeding on protein expression of the lipogenic enzymes FAS, ACLY, and ACC1 with some conflicting results. For FAS protein in mouse liver, reports range from 3-fold downregulation with fasting^72^ to only minor downregulation^53,73,74^ to no change even after 36 hours of starvation^75^. Similarly disparate results have been reported for ACLY and ACC1^53,72,73,76^. Our data indicates that most metabolic enzymes, including the lipogenic proteins, are essentially unchanged when comparing the refed 4h and the starved 24h states despite dramatic regulation of mRNA expression and functional lipogenesis.

One likely explanation for these discrepant results is the definition of “fed” state with some groups using *ad-libitum* conditions, others using refed 6h, others refed 12h, and some using unspecified conditions. The *ad-libitum* condition in particular poses challenges for interpreting results since mice may be in disparate food-saturation states at the time of sacrifice. Pre-fasting mice before refeeding, as many groups have done, creates a more uniform fed condition across biological replicates^32^. Regardless, using clearly defined feeding and fasting conditions will help reconcile studies related to regulation of metabolism. The refeeding period in our data is relatively short (4h) and follows a 16h pre-fast, which may possibly explain the paucity of protein changes between the fed and starved states. Our conditions nonetheless provide three insights into liver metabolic response: **1)** Acute response to feeding (<4h) is characterized by significant upregulation of functional lipogenesis entirely independent of changes in lipogenic enzyme abundance. **2)** mRNA changes of lipogenic transcripts during this phase are dramatic but do not reflect changes in protein. **3)** Abundance of certain proteins like MIG12, PDK4, and PDCD4 (in addition to SREBP-1) are rapidly regulated during feeding, suggesting these early-responder proteins may play important regulatory roles that might be missed under conditions with longer re-feeding windows such as 6h or 12h. Mechanistically, PDCD4 (type I discordant) may be lower in the fed state because of rapid insulin- and S6K-mediated proteasomal degradation as previously reported^77^, which is consistent with the rapid change independent of mRNA change between feeding and starvation.

It is known that lipogenesis is regulated by multiple factors including allosteric metabolites, covalent phosphorylation-dephosphorylation, and changes in enzyme abundance^78^. Allosteric regulators include citrate, which induces the polymerization and activation of ACC1, and fructose-6-phosphate, which activates ACLY^79^. It is well-established that ACC1 activity requires oligomerization of ACC1 subunits in order to convert acetyl-CoA to malonyl-CoA, the immediate substrate required for fatty acid synthesis by FAS^80^. Horton’s group have shown that MIG12 is also a potent activator of ACC1 polymerization^53^, functioning as a non-metabolite allosteric regulator. Interestingly, in our data, *Mid1ip1* / MIG12 displays rapid and concordant upregulation in the fed state and decreases in the starved state. The activity of lipogenic enzymes can also be modulated through phosphorylation of their residues by Akt (ACLY on Ser-455)^79,81^, mTOR (FAS)^82^, AMPK or PKA (ACC on Ser-79 and Ser-1200 respectively)^83–85^. While it lies beyond the scope of the present paper, it is important to continue investigating the physiological significance of site-specific phosphorylation on regulation of lipogenic enzymes *in vivo*.

While several lipogenic regulatory mechanisms are known, the timing of regulation by protein abundance *vs* allosteric/covalent control is not well established. Our data clearly demonstrate that the activation of *de novo* lipogenesis even multiple hours after the onset of feeding is independent of change in the abundance of *bona fide* lipogenic enzymes. Our data shows that MIG12 (fed), PDK4 (starved), and PDCD4 (starved) are some of the few proteins rapidly regulated in response to feeding and starvation. The widespread mRNA-protein-activity discordances we observed emphasize the importance of avoiding conclusions about *de novo* lipogenic activity based solely on protein or mRNA levels.

These findings also beg the question: if the mRNAs for lipogenic genes are markedly induced in the fed state (i.e. *Fasn* ∼40-fold), yet there is no apparent change in protein levels, what is the role for such a large mRNA induction? Recent studies have observed that many proteins, including lipogenic enzymes, are very stable *in vivo* with half-lives greater than 60h^86^. Thus, one possibility is that the rate of translation is relatively slow, so that there is a temporal delay greater than the time frame of acute feeding used in this study. Alternatively, and potentially more interestingly, is the possibility that these mRNAs have an additional role outside of being templates for translation and may function as allosteric regulators, scaffolds, or in the generation of molecular condensates^87,88^. Although beyond the scope of this study, these possible mechanisms are now primed for future investigations.

## MATERIALS AND METHODS

### Animals

All studies followed guidelines approved and in agreement with the Albert Einstein College of Medicine Institutional Animal Care and Use Committee. All experiments were performed on 10-14 week-old C57BL/6J WT male mice from The Jackson Laboratory (#000664). Mice had free access to low fat (4.5%) laboratory mouse chow (LabDiet #5053) and water unless otherwise noted. All mice were group-housed in a facility equipped with a 12h light/dark cycle (7:00am (ZT-0) - 7:00pm (ZT-12)). Prior to fasting and feeding experiments, mice were conditioned to a daylight time-restricted feeding protocol. To initiate conditioning, mice were fasted for 16h by removing chow at 5:00pm (ZT-10 +0) until 9:00am the following day (ZT-2 +1). Mice were then provided with chow for 8h from 9:00am (ZT-2 +1) - 5:00pm (ZT-10 +1). This conditioning cycle was repeated for 2 days. On the day of sample collection, mice were provided chow and 20% sucrose water (g/ml) for 4h from 9:00am (ZT-2 +3) - 1:00pm (ZT-6 +3). Mice sacrificed at 1:00pm (ZT-6 +3) represented the fed condition. For the remaining mice, chow was removed and 20% sucrose water was replaced with regular water at 1:00pm (ZT-6 +3). Mice were then fasted for 24h from 1:00pm (ZT-6 +3) - 1:00pm (ZT-6 +4). Mice sacrificed at 1:00pm (ZT-6 +4) represented the starved condition. Whenever removing chow, mice were transferred to a clean cage to ensure no food particles remained.

### Sample independence and data sourcing

Four independent sets of mice were used to generate data: 24 mice for hepatocyte RNA-seq, 24 mice for hepatocyte proteomics, 10 mice for lipogenesis experiments, and 6 mice for whole liver western blot and RT-qPCR. Male hepatocyte RNA-seq data was previously published by our group (GSE263415) and has been re-analyzed for this paper.

### Whole liver collection

Mice were anesthetized with isoflurane and euthanized by cervical dislocation. Livers were surgically removed, separated from gallbladder, and briefly rinsed with PBS before freeze-clamping with Wollenberger tongs pre-chilled in liquid nitrogen. Frozen livers were homogenized into powder with mortar and pestle pre-chilled with liquid nitrogen and stored at −80°C for downstream processing.

### Pericentral / Periportal Hepatocyte Isolation

Mice were anesthetized with isoflurane on a surgical stage under a heat lamp, and their abdomens were opened. The portal vein (PV) and the lower inferior vena cava (IVC) inferior to the liver were catheterized with 24-gauge catheters (TERUMO, #SR-OX2419CA). For pericentral isolation, the lower IVC catheter was connected to a perfusion line filled with pre-warmed (42°C) 1xPBS with a flow rate of 5 mL/min to wash out blood from the liver through the unconnected PV catheter. While the liver was being perfused with 1xPBS, the diaphragm was cut open and an artery clip was placed on the IVC superior to the liver. Once the liver was pale, the perfusion pump was switched off and the inlet side of the line was switched from PBS to pre-warmed (42°C) enzyme buffer solution (EBS) (15 mL) containing liberase (0.05 mg/mL = 0.26 units/mL) (Sigma-Aldrich, #5401127001). Instructions for making EBS can be found elsewhere^89^. While the perfusion pump was still off, 833 μL of 5-10 mM digitonin (Sigma-Aldrich, #D141-500MG) in 1xPBS in a 1 mL syringe was slowly injected into the unconnected PV catheter over a 10 second interval (∼5 mL/min). Once the digitonin solution was injected, the portal vein was cut and the perfusion pump was immediately restarted from the lower IVC until 15 mL of EBS / liberase was consumed. For periportal isolation, all steps were the same except the perfusion line was connected to the PV catheter and digitonin was injected into the lower IVC catheter. After perfusion was complete, the liver was cut from the mouse and placed in a pre-chilled 10 cm dish containing 20 mL of Krebs-Henseleit (KH) buffer (118 mM NaCl, 4.7 mM KCl, 1.2 mM MgSO_4_, 25 mM NaHCO_3_, 1.2 mM KH_2_PO_4_, pH = 7.4). The liver was briefly minced and shaken in the dish with forceps to release cells. Dissociated liver cells were transferred to a 50 mL conical tube with a 100μm cell strainer (Fisher Scientific, #22363549). Liver cells were centrifuged at 40 x g for 5 min at 4 °C to pellet high density hepatocytes. Supernatant was aspirated, and the pellet (enriched with hepatocytes) was resuspended in 10 mL cold KH buffer. 10 mL of Percoll solution (9 ml Percoll (Sigma-Aldrich, #P4937-500ML) mixed with 1 ml 10× PBS (Invitrogen, #AM9625)) was added to the resuspended cells (20 mL total) and centrifuged at 80 x g for 10 min at 4°C. The supernatant containing dead cells was aspirated, and the pellet was resuspended in 10 mL of KH buffer before a third centrifugation step at 40 x g for 5 min at 4°C. Cells were counted by hemocytometer and viability was assessed using trypan blue staining before snap freezing in liquid nitrogen for downstream processing. Cells for proteomics received one extra wash with 150 mM ammonium acetate in water before snap freezing.

### *De-novo* Palmitate Measurements

Mice were subjected to the time-restricted feeding protocol as described above. For fed condition, mice received intraperitoneal (IP) injection at ZT-0 of 0.9% NaCl in deuterated water (g/mL) (Cambridge Isotope Laboratories, #DLM-4-99) at 30 μl/g body weight, and drinking water was replaced with 6% deuterated water. Two hours after IP at ZT-2, mice were provided with chow and 20% sucrose in 6% deuterated water for four hours. At ZT-6, fed mice were sacrificed and whole liver was collected and processed as described above. Plasma was also collected in EDTA tubes (Sarstedt, #20.1339.100). For starvation condition, mice were fed with chow and 20% sucrose water (no deuterated water) for 4h from ZT-2 to ZT-6 as before. At ZT-6, food was removed and sucrose water was replaced with regular drinking water. The following day at ZT-0, mice received IP injection of 0.9% NaCl in deuterated water at 30 μl/g body weight and regular drinking water was replaced with 6% deuterated water. Starved mice were sacrificed six hours after IP (24 hours after food removal) at ZT-6.

Approximately 30 mg of homogenized liver powder was combined with 300 μl 1 M NaOH in 70% ethanol in water and 20 μl of C17:0 internal standard in 2mM methanol. Approximately 10 ceramic beads (BioSpec Products, #11079110Z) were added, and samples were homogenized at 4°C with BeadBlaster machine (Benchmark Scientific, #D2400) for 2 rounds (30s on / 30s off) at 6 M/S. Samples were incubated at RT for 3h for lipid saponification. Saponified samples were extracted by adding 500 μl of chloroform (Honeywell Riedel-de Haën, #650471), vortexing for 1 min, and centrifuging at 14,000 xg for 10min at RT. 100 μl of bottom chloroform layer was transferred to a new vial and dried under gentle air flow for 30 min. Dried samples were derivatized by adding 50 μl pyridine and 50 μl BSTFA (with 1% TMCS). After heating at 65 °C for 1 hour, the samples (0.2uL in splitless mode) were injected into GC-MS (Agilent) with DB-5MS column. The injection was set to 260 °C, and the oven program was from 100 °C to 310 °C in 22 minutes. Helium was used as carrier gas at the consistent flow rate of 1ml/min. The mass spectrometer was set to full scan mode. For the *de-novo* lipogenesis, the m/z of 313 was used to calculate the palmitate deuterium enrichment. The enrichment was calculated with the area of M0, M1, and M2 of the corresponding fragment with the correction method reported by Jennings and Matthews^90^. The total labeled fraction of palmitate was calculated with fraction enrichment of M1*1+M2*2+M3*3+M4*4. The total palmitate was calculated relative to the concentration of the internal standard (heptadecanoic acid). The newly synthesized palmitate was calculated with the function: Newly synthesized palmitate = Total palmitate * total fraction of labeled palmitate / deuterium enrichment in plasma / tissue weight.

### RNA Isolation and Quantitative PCR (RT-qPCR)

Approximately 200K hepatocytes or 20 mg of homogenized whole liver powder was lysed with 500 μl TRIzol (ThermoFisher, #15596026) and mixed with 100 μl of chloroform (Fisher Scientific, #C2984) before centrifugation at 18,000xg for 20 min at room temperature. Supernatant was transferred to a new tube and mixed with equal volume of 100% ethanol (Sigma-Aldrich, #E7023). Samples were then transferred to RNeasy Mini spin columns (Qiagen, #74106) and washed / eluted according to manufacturer’s protocol. Isolated RNA was quantified by NanoDrop (ThermoFisher Scientific) and used for cDNA strand synthesis with SuperScript IV VILO cDNA synthesis kit (Invitrogen, #11756050). RT-qPCR was performed with PowerUp SYBR Green Master Mix (Applied Biosystems, #A25778) on QuantStudio3 (Applied Biosystems) according to the manufacture’s protocol. Samples were normalized to *Apoa2* gene expression using the ΔΔCt method to determine relative mRNA levels. Primer sequences used in all experiments are listed here: *Fasn* (Forward: GGTCGTTTCTCCATTAAATTCTCAT; Reverse: CTAGAAACTTTCCCAGAAATCTTCC), *Pdcd4* (Forward: GCGGTTAGAAGTGGAGTTGCTG; Reverse: CCACATCATACACCTGTCCAGG), *Mid1ip1* (Forward: GGAGATAGACGAGGTCAGCG; Reverse: ATGGACTTGAGCAGCACGTA), *Apoa2* (Forward: GACGGACCGGATATGCAGAG; Reverse: CTGACCTGACAAGGGGTGTC).

### Bulk RNA-Sequencing

2000 ng of isolated RNA was sent to Azenta Life Sciences for genome-wide mRNA sequencing using NEBNext Ultra RNA Library Prep Kit (New England Biolabs, Ipswich, MA) for NovaSeq (Illumina) sequencing by synthesis. All downstream analysis was performed using R (v4.1.3). Transcript-level raw counts from Kallisto (v0.46.0) were imported into R using the Bioconductor package tximport (v1.20.0), and the expression levels of each gene were estimated. Differential expression analysis between the two groups was performed using the DESeq2 (v1.32.0), which generated log2(fold change) and adjusted P-value.

### Immunoblotting

Commercially available primary antibodies were purchased for the detection of FAS (Abcam, #ab22759, 1:3000, 0.33 µg/mL), PDCD4 (Cell Signaling, #9535, 1:1000, 0.55 µg/mL), and Vinculin (Santa Cruz Biotechnology, #sc-73614, 1:1000, 0.2 µg/mL). MID1IP1 / MIG12 rabbit polyclonal antibody (397E) (1:1000, 4.0 µg/mL) was produced and provided by Dr. Chaiwan Kim and Dr. Jay D. Horton from UT Southwestern Medical Center. 30 mg of homogenized whole liver powder was lysed with RIPA lysis buffer (ThermoFisher Scientific, #89900) containing protease (Sigma-Aldrich, #S8830-20TAB) and phosphatase inhibitors (Sigma-Aldrich, #P5726) by rocking for 10 min at 4°C. After centrifugation at 14,000xg for 10 min at 4°C, total protein in the supernatant was collected, quantified by BCA assay (ThermoFisher Scientific, #23225), and normalized across samples. Normalized protein was mixed with Laemmli buffer (BIO-RAD, #1610747), 2-Mercaptoethanol (BIO-RAD, #1610710) (final concentration 2.5%), and dithiothreitol (DTT) (final concentration 100 mM) and heated at 100 °C for 7 min. 30 µg of protein was loaded per lane in an SDS polyacrylamide gel (BIO-RAD, #4561094) along with molecular weight ladder (BIO-RAD, #1610373). Gel was run at 100V for 1.5 hr with Tris/Glycine/SDS buffer (BIO-RAD, #1610732) and then electrophoretically transferred to polyvinylidene difluoride (PVDF) membrane (Millipore-Sigma, #IPVH00010) at 30V overnight with Tris/Glycine buffer (BIO-RAD, #1610734). Membrane was stained with Ponceau S solution (Millipore-Sigma, P7170) and scanned to assess total protein loading. Membrane was blocked with LICOR blocking buffer (LICORbio, #927-80001) containing 5% milk powder (g/ml) (BIO-RAD, #1706404) for 1 hr at RT and incubated with primary antibody in LICOR blocking buffer (no milk) containing 0.2% Tween-20 overnight at 4°C. Membrane was washed in TBS with Tween20 (TBST) and incubated with horseradish peroxidase-conjugated secondary antibody (Invitrogen, #31460/31437, 1:100,000) for 1 hr at RT in LICOR blocking buffer (no milk) containing 0.2% Tween-20 and 0.02% SDS. Membranes were washed with TBST and visualized by enhanced chemiluminescence (ECL) (Cytiva, #RPN2232) on film (Cytiva, #28906838). ImageJ Gel Analyzer function was used for band quantification.

### Peptide Preparation

Approximately 400K hepatocytes were lysed with RIPA lysis buffer containing protease and phosphatase inhibitors by rocking for 10 min at 4°C as above. After centrifugation at 14,000xg for 10 min at 4°C, total protein in the supernatant was collected, quantified by BCA, and normalized across samples. Normalized lysates were mixed with DTT (final concentration 5 mM) and incubated for 1h at room temperature. Samples were then alkylated with 20 mM iodoacetamide in the dark for 30 min. Afterward, phosphoric acid was added to a final concentration of 1.2%. Samples were diluted in six volumes of binding buffer (90% methanol and 10 mM ammonium bicarbonate, pH 8.0). After gentle mixing, the protein solution was loaded into an S-trap filter (Protifi) and spun at 500 x g for 30 sec. The sample was washed twice with binding buffer. Finally, 1 µg of sequencing grade trypsin (Promega), diluted in 50 mM ammonium bicarbonate, was added into the S-trap filter and samples were digested at 37°C for 18 h. Peptides were eluted in three steps: (i) 40 µl of 50 mM ammonium bicarbonate, (ii) 40 µl of 0.1% TFA and (iii) 40 µl of 60% acetonitrile and 0.1% TFA. The peptide solution was pooled, spun at 1,000 x g for 30 sec and dried in a vacuum centrifuge. Peptide quantification was performed with Pierce Quantification Peptide Assay.

### LC-MS/MS analysis

Samples were acquired using a nanoElute 2 HLPC system (Bruker Daltonik Gmbh) coupled to a timsTOF HT (Bruker Daltonik Gmbh). Peptides were separated with a PepSep XTREME (25 cm x 150 µM, 1.5 µm particle size) with a column temperature of 50°C and through a 20 µm emitter. A 30 minute elution window was used with a 2-35% Solvent B (0.1% formic acid Sigma-Aldrich F0507, acetonitrile) linear gradient at a flow rate of 500 nL/min. The column was then at 95% Solvent B for 5 minutes. The spectra were acquired in dia-PASEF in the range of 100-1700 m/z and 0.6 1/K0 [V-s/cm-2] to 1.6 1/K0 [V-s/cm-2]. Ramp and accumulation times were set to 100. 32 DIA isolation windows were used in the range of 400-1201 m/z and 0.6 1/K0 [V-s/cm-2] to 1.6 1/K0 [V-s/cm-2]. The collision energy was ramped as a function of increasing mobility starting from 20 eV at 0.60 1/K0 to 59 eV at 1.6 1/K0.

### Searching acquired mass spectra

The acquired data was searched using DIA-NN (version 1.9.2). A predicted spectral library was generated in DIA-NN with a SwissProt (2024-01-24 release) mouse database. The maximum number of missed cleaves was set to 1 with trypsin. Cysteine carbamidomethylation was enabled as a fixed modification and methionine oxidation and N-terminal protein acetylation were set as variable modifications. Match between runs was enabled and a false discovery rate of 1% was used. The DIA-NN output was further processed with directLFQ (v0.3.0) to get the ion and protein quantification.

### Proteome informatics

Proteomic data was analyzed in R (4.4.1) with the following packages: biomaRt (2.60.1), dplyr (1.1.4), SummarizedExperiment (1.34.0), ggplot2 (3.5.1), tibble (3.2.1), tidyr (1.3.1), PerformanceAnalytics (2.0.4), MSnbase (2.30.1), reshape2 (1.4.4), limma (3.60.6), UpSetR (1.4.0), ggrepel (0.9.6), plyr (1.8.9), and data.table (1.17.0). A protein was removed from the dataset if it was not detected in all three replicates for at least one condition. Imputation was performed with MinDet imputation^91^ and differential abundance analysis was done with limma^92^. Functional enrichment analysis was performed with Metascape (http://metascape.org) using *M. musculus* as the input and analysis species^93^.

### Gene product identifier mapping and integration

(1) All UniProt accessions from Dia-NN (5592 IDs) were mapped using the UniProt ID Mapping website tool (https://www.uniprot.org/id-mapping) from UniProtKB AC/ID database to Gene Name database. All UniProt mapping results have be uploaded with ID mapping code (Script 2). Results were imported into R and 1:1 mappings were identified (5561 IDs). (2) All Ensembl IDs from tximport (16,728 IDs) were mapped to gene names using the mygene package in R, and 1:1 mappings (16,723 IDs) were identified. (3) Protein ID and mRNA ID sets were merged by gene name, and 5406 IDs were found to have identical, case-sensitive gene names. 11,229 genes were only found in the mRNA set, and 155 genes were only found in the protein set. (4) The IDs only found in the protein set were mapped again using the UniProt ID Mapping website tool from UniProtKB AC/ID database to Ensembl database. Protein IDs with 1:1 mapping to Ensembl IDs (129) were merged with the mRNA ID set by Ensembl ID, and 88 genes shared identical Ensembl IDs. This approached enabled mapping of 5494 / 5592 (98.2%) UniProt accessions to Ensembl IDs in a 1:1 manner.

### Additional Software

R (v4.4.1), Excel (v16.16.27), Prism (v10.4.0), Cytoscape (v3.10.3), Ingenuity Pathway Analysis (v10.4.0), JMP Pro (v18.0.2), ImageJ2 (v2.3.0/1.53f), Powerpoint (v16.16.27), Adobe Illustrator (v25.4.1), BioRender, and Desmos Graphing Calculator were used for data processing, analysis, and figure generation.

## Declaration of generative AI use

GPT-4 Turbo was used by the authors for editing and improving text readability during the writing process. The authors reviewed and edited all revised content and take full responsibility for claims in the published article. GPT-4 Turbo was also used for writing and editing code in R for bioinformatics analysis. All code and results were carefully reviewed.

## Data Availability

Raw fastq files were deposited in the National Center for Biotechnology Information Gene Expression Omnibus database and are available under accession numbers: GSE263415 (previously published male data) and GSE291303 (female data). Proteomics raw files are available upon request from the Corresponding Author. Free access and query of all data is available through the Shiny interactive web application (https://shinoda-lab.shinyapps.io/discorda/).

## Code Availability

The R code and required data used for proteome informatics (Script 1), ID mapping (Script 2), and generation of figures (Script 3) is available for download from the Shinoda Lab GitHub page: https://github.com/kosaku-san/Hepatocyte-Dual-Omics

## Acknowledgements

We thank Dr. Jay D. Horton and Dr. Chaiwan Kim (UT Southwestern) for generously sharing their MIG12 antibody. We thank Shiori Okada for perfusion support; Sofia Krylova and Dr. Yan Tang for antibody validation and literature search; and Dr. Alus Michael Xiaoli for technical assistance. This study was supported by NIH grants DK110063 (JEP), DK110426 (KS), T32GM007491 (MH), S10OD030286 (SS), P30DK020541 (Einstein–Mount Sinai Diabetes Center), and DK026687 (New York Obesity Research Center). The Sidoli and Shinoda labs gratefully acknowledge support from the Hevolution Foundation (via American Federation for Aging Research).

---

**Supplemental Fig. 1.**
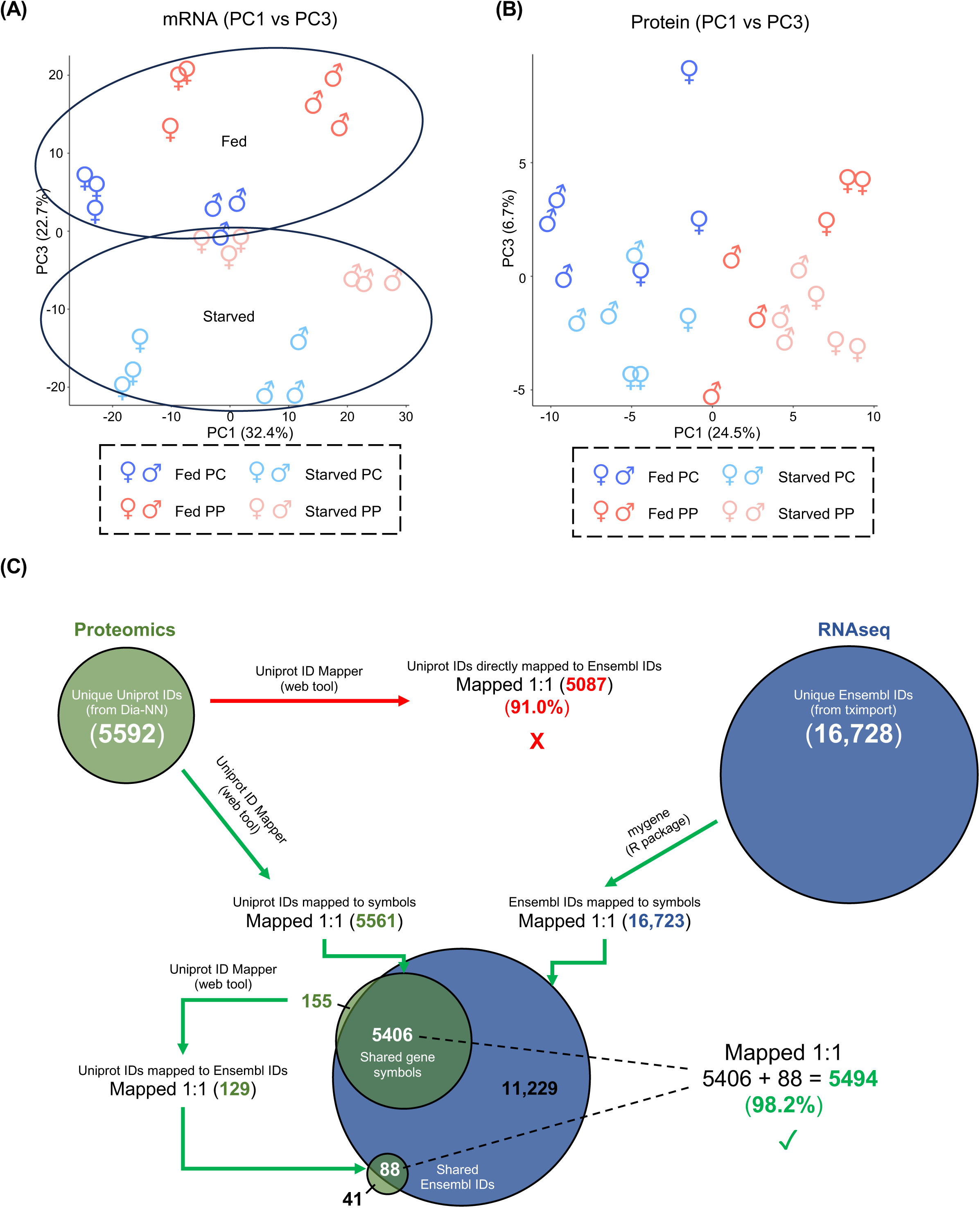
**(A-B)** Principle component analysis (PCA) for **(A)** RNA-seq and **(B)** Proteomics showing principle component 1 (PC1) vs principle component 3 (PC3). Fed and starved samples are circled in mRNA plot. PCAs are based on the top 3% most variable GPs in respective datasets. PCA points represent biological replicates (*n*=3) for all 8 conditions based on coloring shown in the legend. **(C)** Detailed schematic of the method used for mapping protein accession IDs to Ensembl IDs.

**Supplemental Fig. 2.**
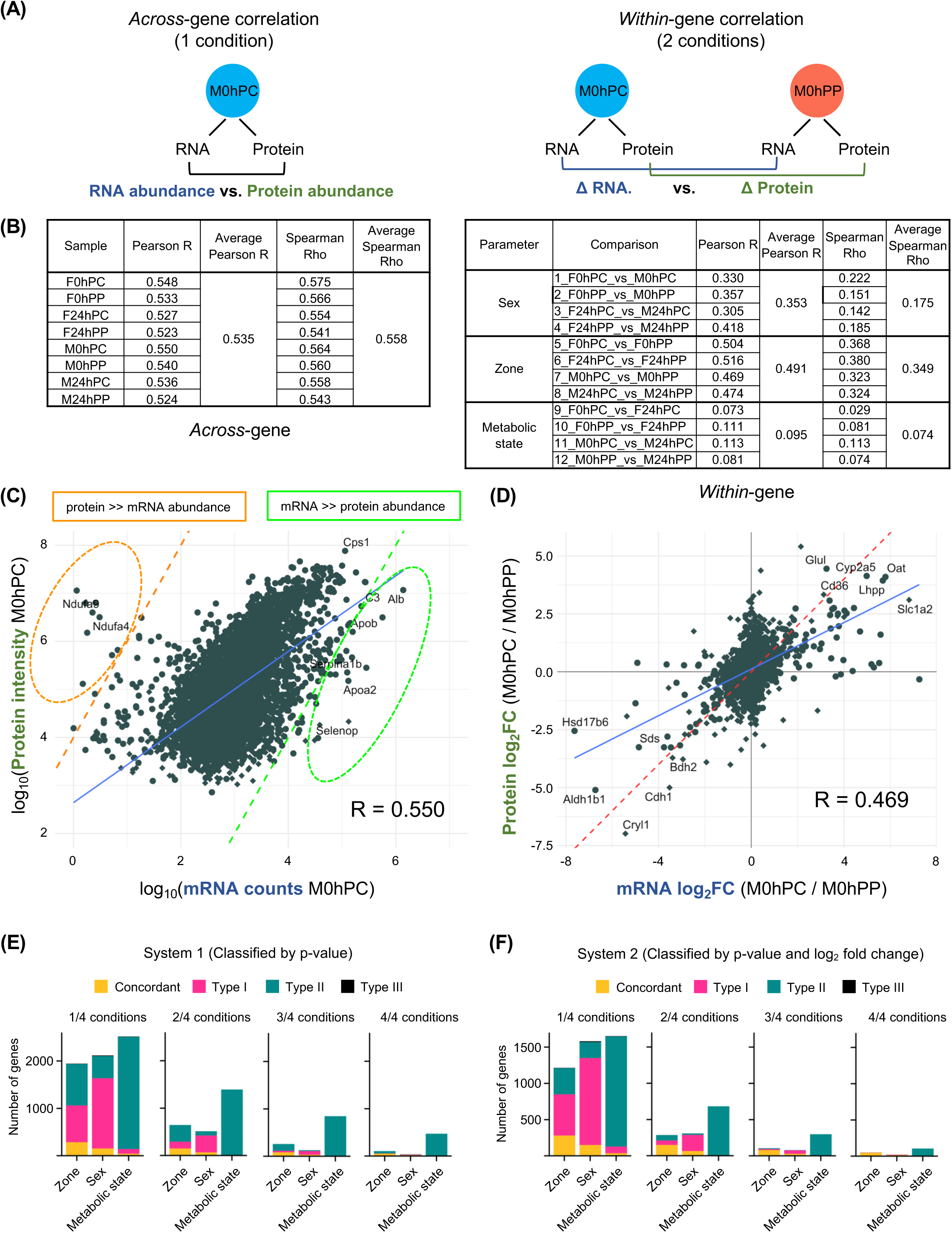
**(A)** Schematic illustrating *across*-gene and *within-*gene correlations. **(B)** Summary table of Pearson *R* and Spearman ρ values for all *across-*gene and *within*-gene comparisons. **(C)** Representative *across*-gene correlation of log_10_ protein intensity vs mRNA intensity for the fed male pericentral condition (“M0hPC”). Orange circle highlights GPs where protein intensity greatly exceeds mRNA intensity. Green circle highlights GPs where mRNA intensity greatly exceeds protein intensity. Dotted orange and green lines represent lines of y = 2x + 4 and y = 2x – 4 respectively. Solid blue line represents linear fit with Pearson *R* shown. Protein and mRNA intensities are based on 3-biological-replicate average. X-axis only shows GPs with log_10_ mRNA intensity > 0. **(D)** Representative *within-*gene correlation scatter plot of protein vs mRNA log_2_ fold change for pericentral (PC) vs periportal (PP) comparison in fed male mice. Dotted red line represents slope of 1. Solid blue line represents linear fit. Pearson *R* is shown. Diamonds represent GPs with imputed protein values. **(E-F)** The number of GPs belonging to the concordant (gold), type I (pink), type II (green), and type III (black) discordant categories as defined by **(E)** adjusted p-value (system 1) or **(F)** fold change with adjusted p-value (system 2) at every cutoff. 1/4 = classified in the corresponding category in one of the four conditions. 2/4 = two of the four conditions. 3/4 = three of four conditions. 4/4 = all four conditions.

**Supplemental Fig. 3.**
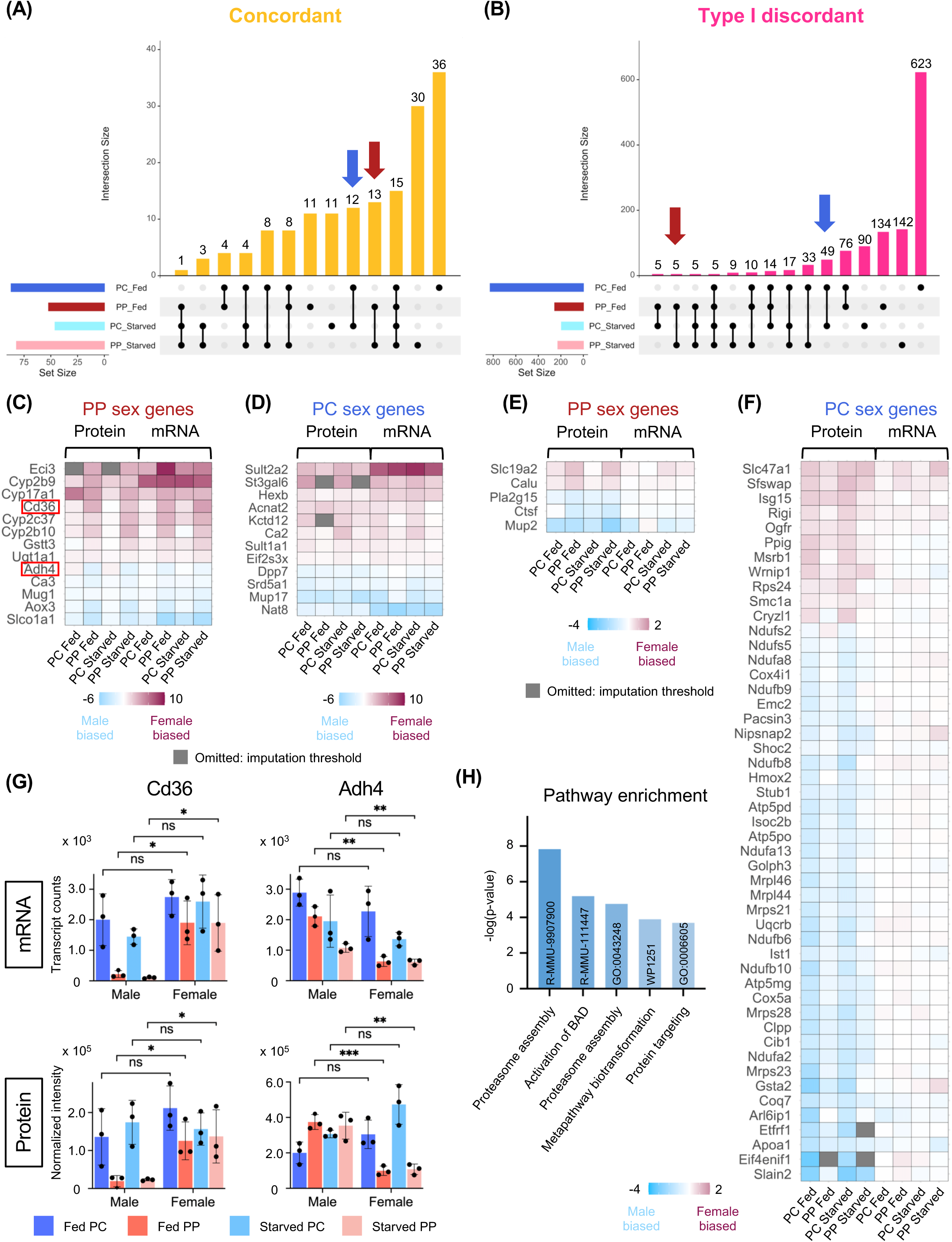
**(A-B)** UpSet plots for **(A)** concordant and **(B)** type I discordant sex-biased GPs with red arrow indicating PP-specific sex GPs and blue arrow indicating PC-specific sex GPs. **(C-D)** Protein and mRNA expression heatmap of **(C)** PP-specific concordant and **(D)** PC-specific concordant sex GPs. **(E-F)** Protein and mRNA expression heatmap of **(E)** PP-specific type I and **(F)** PC-specific type I discordant sex GPs. **(G)** RNA-seq and proteomic values for representative PP-specific concordant sex GPs (*Cd36* and *Adh4)* (*n*=3 mice per condition). Statistical comparisons are based on unpaired Student’s t-test performed in Prism. (n.s. = not significant, *p<0.05, **p<0.01, ***p<0.001). (**H**) Metascape pathway enrichment analysis of type I discordant GPs from the PC fed and PP fed condition (76 GPs total). For **(C-F)**, the proteins shown in gray were omitted from the analysis because they did not meet the criteria for imputation using the MinDet algorithm.

**Supplemental Fig. 4.**
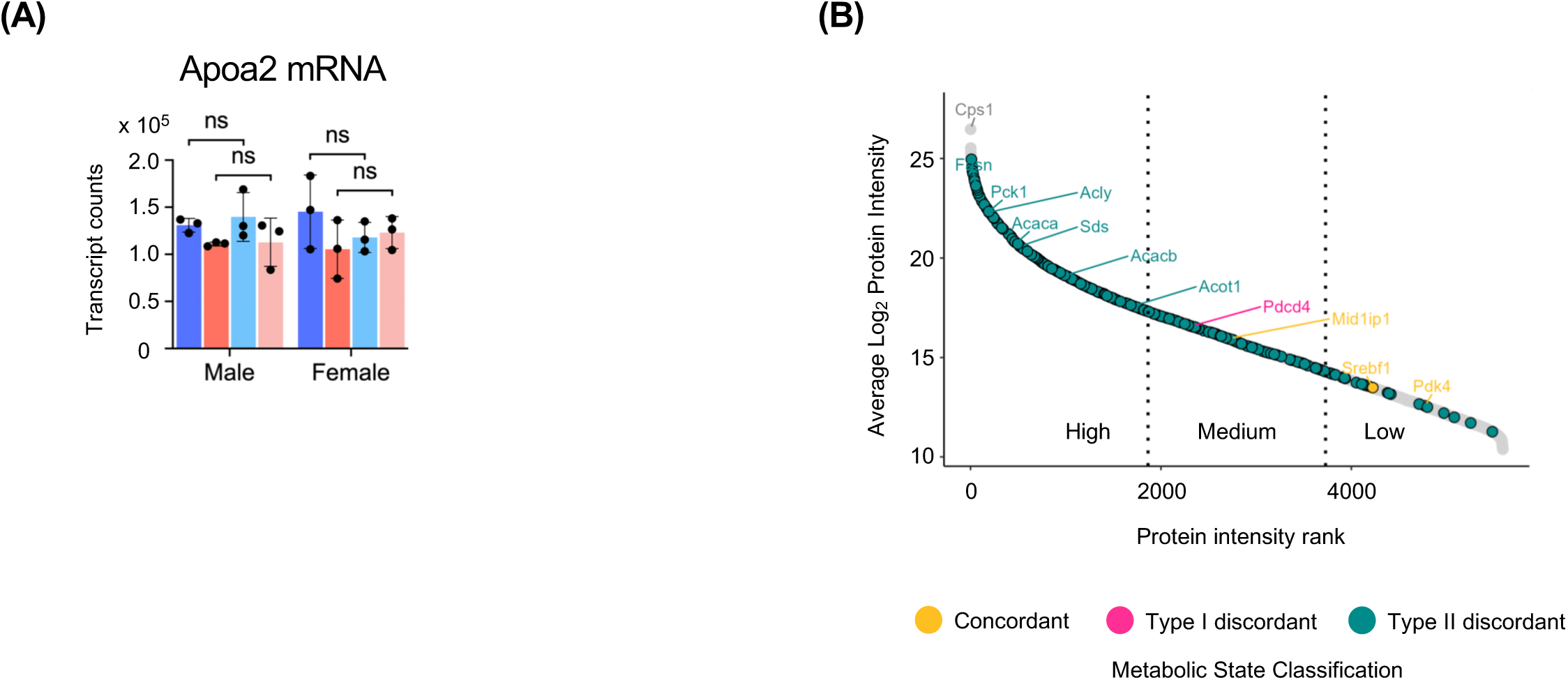
(**A**) RNA-seq transcript counts for the RT-qPCR housekeeping gene, *Apoa2* (*n*=3 mice per condition). Statistical comparisons are based on unpaired Student’s t-test performed in Prism. (n.s. = not significant). (**B**) Plot of log_2_ protein intensity vs protein abundance rank with proteins colored based on classification in the metabolic state parameter. Plot is divided into three regions to separate high, medium, and low abundance proteins.

**Supplemental Table 1:**
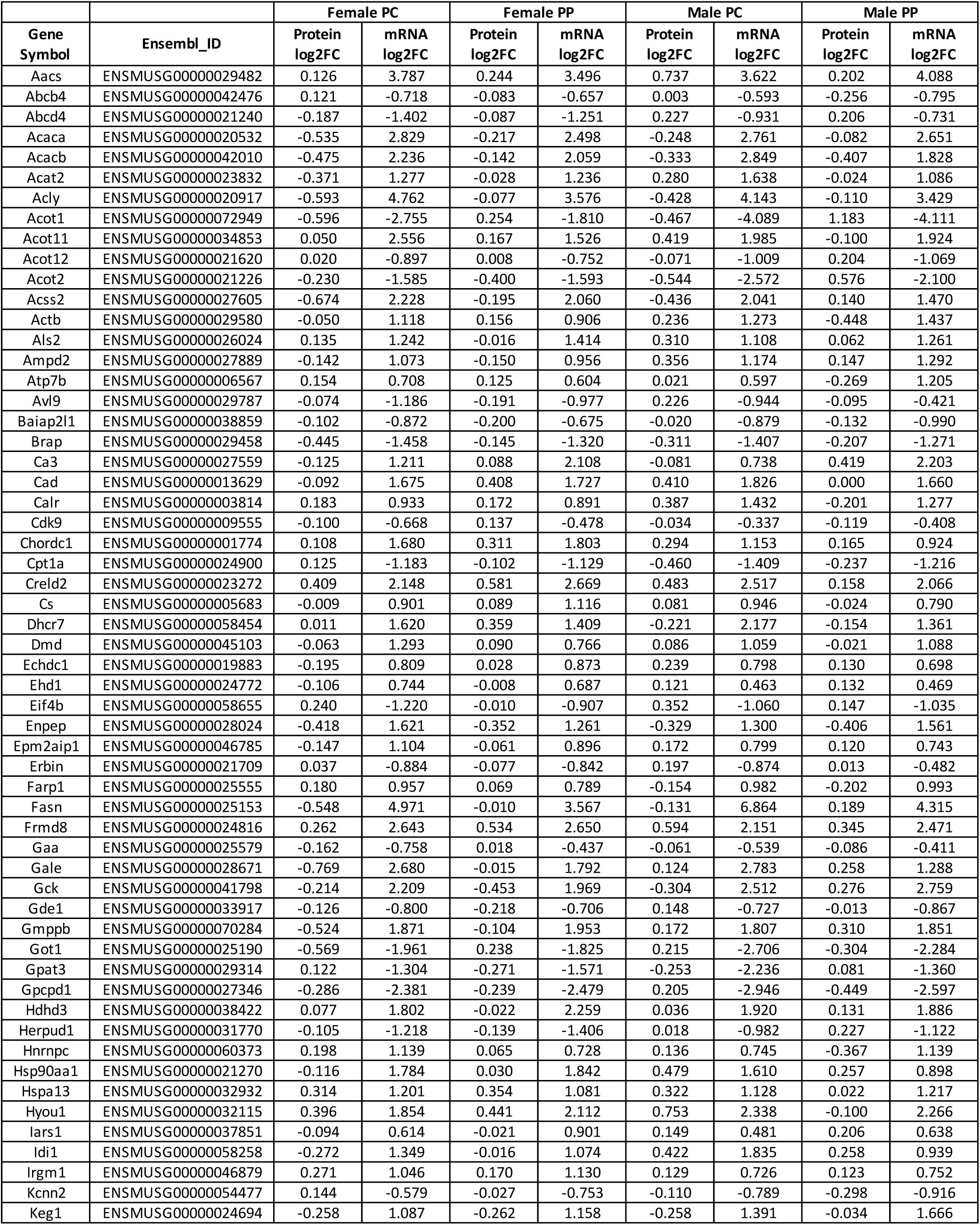
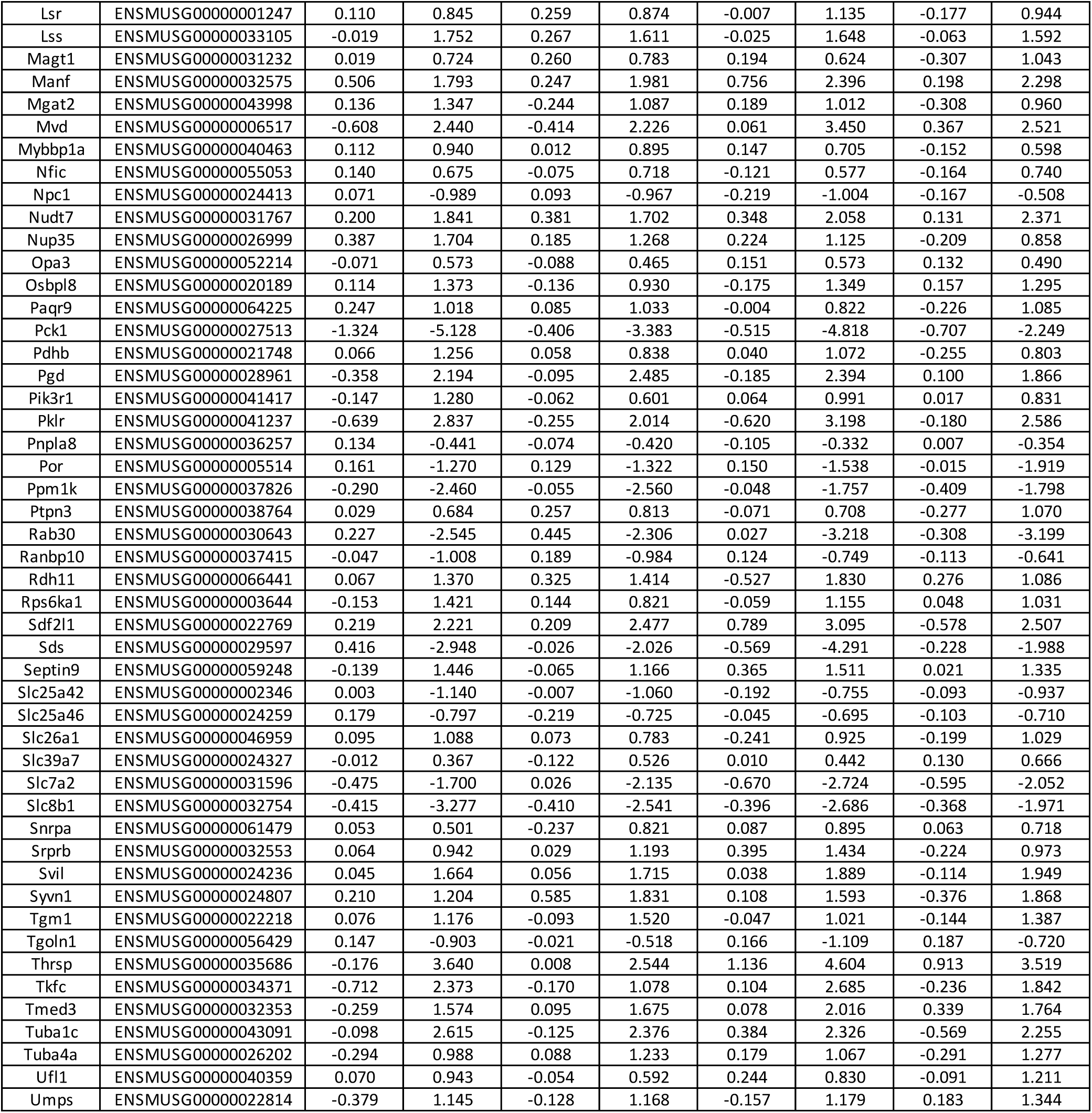
106 “core” type II discordant GPs and their relative changes during the fed-starvation transition.

## REFERENCES

1. Saltiel AR, Kahn CR. Insulin signalling and the regulation of glucose and lipid metabolism. Nature. 2001;414(6865):799–806.

2. Oh KJ, Han HS, Kim MJ, Koo SH. CREB and FoxO1: two transcription factors for the regulation of hepatic gluconeogenesis. BMB Rep. 2013;46(12):567–574.

3. Jitrapakdee S. Transcription factors and coactivators controlling nutrient and hormonal regulation of hepatic gluconeogenesis. Int J Biochem Cell Biol. 2012;44(1):33–45.

4. Lin HV, Accili D. Hormonal regulation of hepatic glucose production in health and disease. Cell Metab. 2011;14(1):9–19.

5. Chen G, Liang G, Ou J, Goldstein JL, Brown MS. Central role for liver X receptor in insulin-mediated activation of Srebp-1c transcription and stimulation of fatty acid synthesis in liver. Proc Natl Acad Sci U S A. 2004;101(31):11245–11250.

6. Bideyan L, Nagari R, Tontonoz P. Hepatic transcriptional responses to fasting and feeding. Genes Dev. 2021;35(9-10):635–657.

7. Solinas G, Boren J, Dulloo AG. De novo lipogenesis in metabolic homeostasis: More friend than foe? Mol Metab. 2015;4(5):367–377.

8. Ameer F, Scandiuzzi L, Hasnain S, Kalbacher H, Zaidi N. De novo lipogenesis in health and disease. Metabolism. 2014;63(7):895–902.

9. Lonardo A, Nascimbeni F, Ballestri S, et al. Sex Differences in Nonalcoholic Fatty Liver Disease: State of the Art and Identification of Research Gaps. Hepatology. 2019;70(4):1457–1469.

10. Waxman DJ, Holloway MG. Sex differences in the expression of hepatic drug metabolizing enzymes. Mol Pharmacol. 2009;76(2):215–228.

11. Smiriglia A, Lorito N, Serra M, Perra A, Morandi A, Kowalik MA. Sex difference in liver diseases: How preclinical models help to dissect the sex-related mechanisms sustaining NAFLD and hepatocellular carcinoma. iScience. 2023;26(12):108363.

12. Clodfelter KH, Holloway MG, Hodor P, Park SH, Ray WJ, Waxman DJ. Sex-dependent liver gene expression is extensive and largely dependent upon signal transducer and activator of transcription 5b (STAT5b): STAT5b-dependent activation of male genes and repression of female genes revealed by microarray analysis. Mol Endocrinol. 2006;20(6):1333–1351.

13. Zhang Y, Klein K, Sugathan A, et al. Transcriptional profiling of human liver identifies sex-biased genes associated with polygenic dyslipidemia and coronary artery disease. PLoS One. 2011;6(8):e23506.

14. Lau-Corona D, Ma H, Vergato C, et al. Constitutively Active STAT5b Feminizes Mouse Liver Gene Expression. Endocrinology. 2022;163(5).

15. Pincus SM, Gevers EF, Robinson IC, et al. Females secrete growth hormone with more process irregularity than males in both humans and rats. Am J Physiol. 1996;270(1 Pt 1):E107–115.

16. Ben-Moshe S, Itzkovitz S. Spatial heterogeneity in the mammalian liver. Nat Rev Gastroenterol Hepatol. 2019;16(7):395–410.

17. de Godoy LM, Olsen JV, Cox J, et al. Comprehensive mass-spectrometry-based proteome quantification of haploid versus diploid yeast. Nature. 2008;455(7217):1251–1254.

18. Newman JR, Ghaemmaghami S, Ihmels J, et al. Single-cell proteomic analysis of S. cerevisiae reveals the architecture of biological noise. Nature. 2006;441(7095):840–846.

19. Ghaemmaghami S, Huh WK, Bower K, et al. Global analysis of protein expression in yeast. Nature. 2003;425(6959):737–741.

20. Lu R, Markowetz F, Unwin RD, et al. Systems-level dynamic analyses of fate change in murine embryonic stem cells. Nature. 2009;462(7271):358–362.

21. Schwanhausser B, Busse D, Li N, et al. Global quantification of mammalian gene expression control. Nature. 2011;473(7347):337–342.

22. Chi Y, Youn DY, Xiaoli AM, et al. Comparative impact of dietary carbohydrates on the liver transcriptome in two strains of mice. Physiol Genomics. 2021;53(11):456–472.

23. Chi Y, Youn DY, Xiaoli AM, et al. Regulation of gene expression during the fasting-feeding cycle of the liver displays mouse strain specificity. J Biol Chem. 2020;295(15):4809–4821.

24. Deaciuc IV, Song Z, Peng X, et al. Genome-wide transcriptome expression in the liver of a mouse model of high carbohydrate diet-induced liver steatosis and its significance for the disease. Hepatol Int. 2008;2(1):39–49.

25. Renaud HJ, Cui JY, Lu H, Klaassen CD. Effect of diet on expression of genes involved in lipid metabolism, oxidative stress, and inflammation in mouse liver-insights into mechanisms of hepatic steatosis. PLoS One. 2014;9(2):e88584.

26. Teufel A, Itzel T, Erhart W, et al. Comparison of Gene Expression Patterns Between Mouse Models of Nonalcoholic Fatty Liver Disease and Liver Tissues From Patients. Gastroenterology. 2016;151(3):513–525 e510.

27. Softic S, Gupta MK, Wang GX, et al. Divergent effects of glucose and fructose on hepatic lipogenesis and insulin signaling. J Clin Invest. 2017;127(11):4059–4074.

28. Bishop DJ, Hoffman NJ, Taylor DF, Saner NJ, Lee MJ, Hawley JA. Discordant skeletal muscle gene and protein responses to exercise. Trends Biochem Sci. 2023;48(11):927–936.

29. Makhnovskii PA, Zgoda VG, Bokov RO, et al. Regulation of Proteins in Human Skeletal Muscle: The Role of Transcription. Sci Rep. 2020;10(1):3514.

30. Youn DY, Xiaoli AM, Kwon H, Yang F, Pessin JE. The subunit assembly state of the Mediator complex is nutrient-regulated and is dysregulated in a genetic model of insulin resistance and obesity. J Biol Chem. 2019;294(23):9076–9083.

31. Youn DY, Xiaoli AM, Zong H, et al. The Mediator complex kinase module is necessary for fructose regulation of liver glycogen levels through induction of glucose-6-phosphatase catalytic subunit (G6pc). Mol Metab. 2021;48:101227.

32. Okada J, Landgraf A, Xiaoli AM, et al. Spatial hepatocyte plasticity of gluconeogenesis during the metabolic transitions between fed, fasted and starvation states. bioRxiv. 2024.

33. Lindros KO, Penttila KE. Digitonin-collagenase perfusion for efficient separation of periportal or perivenous hepatocytes. Biochem J. 1985;228(3):757–760.

34. Quistorff B. Gluconeogenesis in periportal and perivenous hepatocytes of rat liver, isolated by a new high-yield digitonin/collagenase perfusion technique. Biochem J. 1985;229(1):221–226.

35. Pundir S, Martin MJ, O’Donovan C, UniProt C. UniProt Tools. Curr Protoc Bioinformatics. 2016;53:1 29 21-21 29 15.

36. Ruffier M, Kahari A, Komorowska M, et al. Ensembl core software resources: storage and programmatic access for DNA sequence and genome annotation. Database (Oxford). 2017;2017(1).

37. Buccitelli C, Selbach M. mRNAs, proteins and the emerging principles of gene expression control. Nat Rev Genet. 2020;21(10):630–644.

38. Ben-Moshe S, Shapira Y, Moor AE, et al. Spatial sorting enables comprehensive characterization of liver zonation. Nat Metab. 2019;1(9):899–911.

39. Braeuning A, Ittrich C, Kohle C, et al. Differential gene expression in periportal and perivenous mouse hepatocytes. FEBS J. 2006;273(22):5051–5061.

40. Saito K, Negishi M, James Squires E. Sexual dimorphisms in zonal gene expression in mouse liver. Biochem Biophys Res Commun. 2013;436(4):730–735.

41. Rosenberger FA, Thielert M, Strauss MT, et al. Spatial single-cell mass spectrometry defines zonation of the hepatocyte proteome. Nat Methods. 2023;20(10):1530–1536.

42. Droin C, Kholtei JE, Bahar Halpern K, et al. Space-time logic of liver gene expression at sub-lobular scale. Nat Metab. 2021;3(1):43–58.

43. Lau-Corona D, Suvorov A, Waxman DJ. Feminization of Male Mouse Liver by Persistent Growth Hormone Stimulation: Activation of Sex-Biased Transcriptional Networks and Dynamic Changes in Chromatin States. Mol Cell Biol. 2017;37(19).

44. Zhang Y, Laz EV, Waxman DJ. Dynamic, sex-differential STAT5 and BCL6 binding to sex-biased, growth hormone-regulated genes in adult mouse liver. Mol Cell Biol. 2012;32(4):880–896.

45. MacLeod JN, Pampori NA, Shapiro BH. Sex differences in the ultradian pattern of plasma growth hormone concentrations in mice. J Endocrinol. 1991;131(3):395–399.

46. Goldfarb CN, Karri K, Pyatkov M, Waxman DJ. Interplay Between GH-regulated, Sex-biased Liver Transcriptome and Hepatic Zonation Revealed by Single-Nucleus RNA Sequencing. Endocrinology. 2022;163(7).

47. Kramer A, Green J, Pollard J, Jr., Tugendreich S. Causal analysis approaches in Ingenuity Pathway Analysis. Bioinformatics. 2014;30(4):523–530.

48. Holloway MG, Miles GD, Dombkowski AA, Waxman DJ. Liver-specific hepatocyte nuclear factor-4alpha deficiency: greater impact on gene expression in male than in female mouse liver. Mol Endocrinol. 2008;22(5):1274–1286.

49. Lamming DW, Mihaylova MM, Katajisto P, et al. Depletion of Rictor, an essential protein component of mTORC2, decreases male lifespan. Aging Cell. 2014;13(5):911–917.

50. Rooney J, Oshida K, Vasani N, et al. Activation of Nrf2 in the liver is associated with stress resistance mediated by suppression of the growth hormone-regulated STAT5b transcription factor. PLoS One. 2018;13(8):e0200004.

51. Shen WK, Chen SY, Gan ZQ, et al. AnimalTFDB 4.0: a comprehensive animal transcription factor database updated with variation and expression annotations. Nucleic Acids Res. 2023;51(D1):D39–D45.

52. Kosters A, Sun D, Wu H, et al. Sexually dimorphic genome-wide binding of retinoid X receptor alpha (RXRalpha) determines male-female differences in the expression of hepatic lipid processing genes in mice. PLoS One. 2013;8(8):e71538.

53. Kim CW, Moon YA, Park SW, Cheng D, Kwon HJ, Horton JD. Induced polymerization of mammalian acetyl-CoA carboxylase by MIG12 provides a tertiary level of regulation of fatty acid synthesis. Proc Natl Acad Sci U S A. 2010;107(21):9626–9631.

54. Park BY, Jeon JH, Go Y, et al. PDK4 Deficiency Suppresses Hepatic Glucagon Signaling by Decreasing cAMP Levels. Diabetes. 2018;67(10):2054–2068.

55. Brito Querido J, Sokabe M, Diaz-Lopez I, et al. Human tumor suppressor protein Pdcd4 binds at the mRNA entry channel in the 40S small ribosomal subunit. Nat Commun. 2024;15(1):6633.

56. Ye X, Huang Z, Li Y, et al. Human tumor suppressor PDCD4 directly interacts with ribosomes to repress translation. Cell Res. 2024;34(7):522–525.

57. Hatchwell L, Harney DJ, Cielesh M, et al. Multi-omics Analysis of the Intermittent Fasting Response in Mice Identifies an Unexpected Role for HNF4alpha. Cell Rep. 2020;30(10):3566–3582 e3564.

58. Yu Y, Jiang L, Wang H, et al. Hepatic transferrin plays a role in systemic iron homeostasis and liver ferroptosis. Blood. 2020;136(6):726–739.

59. Luo QQ, Zhou G, Huang SN, Mu MD, Chen YJ, Qian ZM. Ghrelin is Negatively Correlated with Iron in the Serum in Human and Mice. Ann Nutr Metab. 2018;72(1):37–42.

60. Tian J, Wu J, Chen X, et al. BHLHE40, a third transcription factor required for insulin induction of SREBP-1c mRNA in rodent liver. Elife. 2018;7.

61. Li J, Takaishi K, Cook W, McCorkle SK, Unger RH. Insig-1 “brakes” lipogenesis in adipocytes and inhibits differentiation of preadipocytes. Proc Natl Acad Sci U S A. 2003;100(16):9476–9481.

62. Yook JS, Taxin ZH, Yuan B, et al. The SLC25A47 locus controls gluconeogenesis and energy expenditure. Proc Natl Acad Sci U S A. 2023;120(9):e2216810120.

63. Goldberg D, Buchshtab N, Charni-Natan M, Goldstein I. Transcriptional cascades during fasting amplify gluconeogenesis and instigate a secondary wave of ketogenic gene transcription. Liver Int. 2024;44(11):2964–2982.

64. Shimomura I, Bashmakov Y, Ikemoto S, Horton JD, Brown MS, Goldstein JL. Insulin selectively increases SREBP-1c mRNA in the livers of rats with streptozotocin-induced diabetes. Proc Natl Acad Sci U S A. 1999;96(24):13656–13661.

65. Softic S, Cohen DE, Kahn CR. Role of Dietary Fructose and Hepatic De Novo Lipogenesis in Fatty Liver Disease. Dig Dis Sci. 2016;61(5):1282–1293.

66. Fu X, Fletcher JA, Deja S, Inigo-Vollmer M, Burgess SC, Browning JD. Persistent fasting lipogenesis links impaired ketogenesis with citrate synthesis in humans with nonalcoholic fatty liver. J Clin Invest. 2023;133(9).

67. Fu X, Deja S, Fletcher JA, et al. Measurement of lipogenic flux by deuterium resolved mass spectrometry. Nat Commun. 2021;12(1):3756.

68. Zhao J, Zhai B, Gygi SP, Goldberg AL. mTOR inhibition activates overall protein degradation by the ubiquitin proteasome system as well as by autophagy. Proc Natl Acad Sci U S A. 2015;112(52):15790–15797.

69. Gao D, Wan L, Inuzuka H, et al. Rictor forms a complex with Cullin-1 to promote SGK1 ubiquitination and destruction. Mol Cell. 2010;39(5):797–808.

70. Kim SJ, DeStefano MA, Oh WJ, et al. mTOR complex 2 regulates proper turnover of insulin receptor substrate-1 via the ubiquitin ligase subunit Fbw8. Mol Cell. 2012;48(6):875–887.

71. Killinger BJ, Petyuk VA, Wright AT. Detecting differential protein abundance by combining peptide level P-values. Mol Omics. 2020;16(6):554–562.

72. Zeng H, Qin H, Liao M, et al. CD36 promotes de novo lipogenesis in hepatocytes through INSIG2-dependent SREBP1 processing. Mol Metab. 2022;57:101428.

73. Liu B, Mao X, Huang D, Li F, Dong N. Novel role of NLRP3-inflammasome in regulation of lipogenesis in fasting-induced hepatic steatosis. Diabetes Metab Syndr Obes. 2019;12:801–811.

74. Kristensen CM, Jessen H, Ringholm S, Pilegaard H. Muscle PGC-1alpha in exercise and fasting-induced regulation of hepatic UPR in mice. Acta Physiol (Oxf). 2018;224(4):e13158.

75. Wei H, Weaver YM, Yang C, et al. Proteolytic activation of fatty acid synthase signals pan-stress resolution. Nat Metab. 2024;6(1):113–126.

76. Guzman M, Bijleveld C, Geelen MJ. Flexibility of zonation of fatty acid oxidation in rat liver. Biochem J. 1995;311 (Pt 3)(Pt 3):853–860.

77. Dorrello NV, Peschiaroli A, Guardavaccaro D, Colburn NH, Sherman NE, Pagano M. S6K1- and betaTRCP-mediated degradation of PDCD4 promotes protein translation and cell growth. Science. 2006;314(5798):467–471.

78. Hillgartner FB, Salati LM, Goodridge AG. Physiological and molecular mechanisms involved in nutritional regulation of fatty acid synthesis. Physiol Rev. 1995;75(1):47–76.

79. Potapova IA, El-Maghrabi MR, Doronin SV, Benjamin WB. Phosphorylation of recombinant human ATP:citrate lyase by cAMP-dependent protein kinase abolishes homotropic allosteric regulation of the enzyme by citrate and increases the enzyme activity. Allosteric activation of ATP:citrate lyase by phosphorylated sugars. Biochemistry. 2000;39(5):1169–1179.

80. Beaty NB, Lane MD. The polymerization of acetyl-CoA carboxylase. J Biol Chem. 1983;258(21):13051–13055.

81. Berwick DC, Hers I, Heesom KJ, Moule SK, Tavare JM. The identification of ATP-citrate lyase as a protein kinase B (Akt) substrate in primary adipocytes. J Biol Chem. 2002;277(37):33895–33900.

82. Jensen-Urstad AP, Song H, Lodhi IJ, et al. Nutrient-dependent phosphorylation channels lipid synthesis to regulate PPARalpha. J Lipid Res. 2013;54(7):1848–1859.

83. Carling D, Zammit VA, Hardie DG. A common bicyclic protein kinase cascade inactivates the regulatory enzymes of fatty acid and cholesterol biosynthesis. FEBS Lett. 1987;223(2):217–222.

84. Ha J, Daniel S, Broyles SS, Kim KH. Critical phosphorylation sites for acetyl-CoA carboxylase activity. J Biol Chem. 1994;269(35):22162–22168.

85. Dyck JR, Kudo N, Barr AJ, Davies SP, Hardie DG, Lopaschuk GD. Phosphorylation control of cardiac acetyl-CoA carboxylase by cAMP-dependent protein kinase and 5’-AMP activated protein kinase. Eur J Biochem. 1999;262(1):184–190.

86. Rolfs Z, Frey BL, Shi X, Kawai Y, Smith LM, Welham NV. An atlas of protein turnover rates in mouse tissues. Nat Commun. 2021;12(1):6778.

87. Castello A, Hentze MW, Preiss T. Metabolic Enzymes Enjoying New Partnerships as RNA-Binding Proteins. Trends Endocrinol Metab. 2015;26(12):746–757.

88. Curtis NJ, Jeffery CJ. The expanding world of metabolic enzymes moonlighting as RNA binding proteins. Biochem Soc Trans. 2021;49(3):1099–1108.

89. Mederacke I, Dapito DH, Affo S, Uchinami H, Schwabe RF. High-yield and high-purity isolation of hepatic stellate cells from normal and fibrotic mouse livers. Nat Protoc. 2015;10(2):305–315.

90. Jennings ME, 2nd, Matthews DE. Determination of complex isotopomer patterns in isotopically labeled compounds by mass spectrometry. Anal Chem. 2005;77(19):6435–6444.

91. Peng H, Wang H, Kong W, Li J, Goh WWB. Optimizing differential expression analysis for proteomics data via high-performing rules and ensemble inference. Nat Commun. 2024;15(1):3922.

92. Ritchie ME, Phipson B, Wu D, et al. limma powers differential expression analyses for RNA-sequencing and microarray studies. Nucleic Acids Res. 2015;43(7):e47.

93. Zhou Y, Zhou B, Pache L, et al. Metascape provides a biologist-oriented resource for the analysis of systems-level datasets. Nat Commun. 2019;10(1):1523.

